# Role of autoregulation and relative synthesis of operon partners in alternative sigma factor networks

**DOI:** 10.1101/032359

**Authors:** Jatin Narula, Abhinav Tiwari, Oleg A. Igoshin

**Author notes:** These authors contributed equally to this work (Jatin Narula, Abhinav Tiwari).

## Abstract

Despite the central role of alternative sigma factors in bacterial stress response and virulence their regulation remains incompletely understood. Here we investigate one of the best-studied examples of alternative sigma factors: the σ^B^ network that controls the general stress response of *Bacillus subtilis* to uncover widely relevant general design principles that describe the structure-function relationship of alternative sigma factor regulatory networks. We show that the relative stoichiometry of the synthesis rates of σ^B^, its anti-sigma factor RsbW and the anti-anti-sigma factor RsbV plays a critical role in shaping the network behavior by forcing the σ^B^ network to function as an ultrasensitive negative feedback loop. We further demonstrate how this negative feedback regulation insulates alternative sigma factor activity from competition with the housekeeping sigma factor for RNA polymerase and allows multiple stress sigma factors to function simultaneously with little competitive interference.

**Major Subject Areas:** Computational and systems biology, Microbiology & Infectious disease

**Research Organism:** *Bacillus subtilis*

## Introduction

Bacteria survive in stressful environmental conditions by inducing dramatic changes in their gene expression patterns [1,2]. For a variety of stresses, these global changes in gene expression are brought about by the activation of alternative σ-factors that bind the RNA polymerase core enzyme and direct it towards the appropriate stress response regulons [3]. Consequently, to ensure that these σ-factors are only active under specific environmental conditions, bacteria have evolved regulatory systems to control their production, activity and availability [3,4]. These regulatory networks can be highly complex but frequently share features such as anti-σ-factors, partner switching mechanisms and proteolytic activation [4]. The complexity of these networks has impeded a clear mechanistic understanding of the resulting dynamical properties. In this study, we focus on one of the best studied examples of alternative σ-factors, the general stress-response regulating σ^B^ in *Bacillus subtilis* [5] to understand how the structure of the σ-factor regulatory networks is related to their functional response.

The σ^B^-mediated response is triggered by diverse energy and environmental stress signals and activates expression of a broad array of genes needed for cell survival in these conditions [5]. Activity of σ^B^ is tightly regulated by a partner-switching network (Fig. 1A) comprising σ^B^, its antagonist anti-σ-factor RsbW, and anti-anti-σ-factor RsbV. In the absence of stress, RsbW dimer (RsbW_2_) binds to σ^B^ and prevents its association with RNA polymerase thereby keeping the σ^B^ regulon OFF. Under these conditions most of RsbV is kept in the phosphorylated form (RsbV~P) by the kinase activity of RsbW_2_. RsbV~P has a low affinity for RsbW_2_ and cannot interact with it effectively [6]. However, in the presence of stress, RsbV~P is dephosphorylated by one or both of the dedicated phosphatase complexes (thereafter, phosphatases): RsbQP for energy stress and RsbTU for environmental stress [7–10]. Dephosphorylated RsbV attacks the σ^B^-RsbW_2_ complex to induce σ^B^ release, thereby turning the σ^B^ regulon ON [11]. Notably, the genes encoding σ^B^ and its regulators lie within a σ^B^-controlled operon [12], thereby resulting in positive and negative feedback loops.

**Figure 1.**
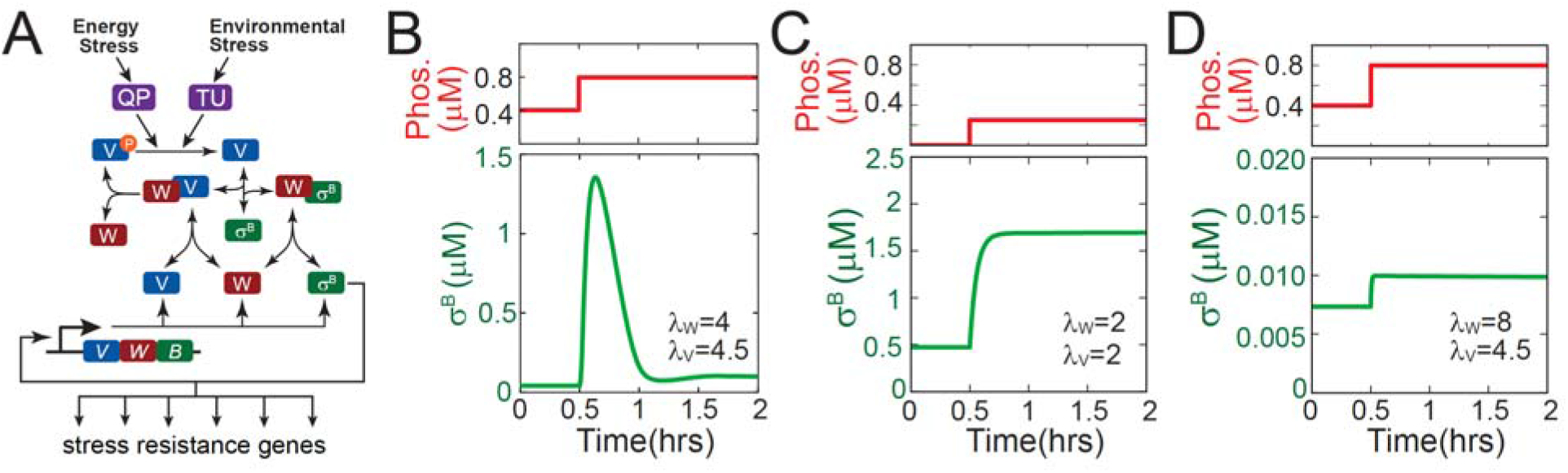
σ^B^ general stress response network. **A.** Network diagram of the σ^B^ general stress response. Energy and environmental stresses activate the stress-sensing phosphatases RsbQP (QP) and RsbTU (TU) which dephosphorylate RsbV which in turn activates σ^B^ by releasing it from the σ^B^-RsbW_2_ complex. Note only the monomeric forms of RsbW and RsbV have been shown for simplicity. **B-D**. Dynamics of free σ^B^ in response to a step-increase in phosphatase concentration for different combinations of the relative synthesis rates of σ^B^ operon partners (λ_W_ = RsbW_T_/B_T_, λ_V_ = RsbV_T_/B_T_).

Recently, it was shown that under energy stress σ^B^ is activated in a stochastic series of transient pulses and increasing stress resulted in higher pulse frequencies [13]. It has also been shown that increase in environmental stressor such as ethanol leads to a single σ^B^ pulse with an amplitude that is sensitive to the rate of stressor increase [14]. While it is clear that the pulsatile activation of σ^B^ is rooted in the complex architecture of its regulatory network (Fig. 1A) its mechanism is not fully understood. Previous mathematical models of the σ^B^ network either did not produce the pulsatile response [15] or made simplifications to the network [13] that are somewhat inconsistent with experimentally observed details. As a result, it remains unclear which design features of the σ^B^ network enable its functional properties.

To address these issues we develop a detailed mathematical model of the σ^B^ network and examine its dynamics to understand the mechanistic principles underlying the pulsatile response. By decoupling the post-translational and transcriptional components of the network we show that an ultrasensitive negative feedback between the two is the basis for σ^B^ pulsing. Moreover we find that the relative synthesis rates of σ^B^ and its operon partners RsbW and RsbV, plays a critical role in determining the nature of the σ^B^ response. We also use our model, together with previously published experimental data from [13,14], to explain how the σ^B^ network is able to encode the rate of stress increase and the size of stochastic bursts of stress phosphatase into the amplitudes of σ^B^ pulses.

We further develop this model to investigate how the network functions in the context of other σ-factors. As in many other bacteria, σ^B^ is one of the many σ-factors that complex with RNA-polymerase core that is present in limited amounts [3,16]. Therefore, when induced these alternative σ-factors compete with one another and the housekeeping σ-factor σ^A^ for RNA polymerase. We use our model to investigate how the design of this network enables it to function even in the presence of competition from σ^A^ which has a significantly higher affinity for RNA polymerase [17]. Lastly, we investigate how multiple alternative σ-factors compete when cells are exposed to multiple stresses simultaneously. Using our model we identify design features that are ubiquitous in stress σ-factor regulation and critical to bacterial survival under diverse types of stresses.

## Results

### Biochemically accurate model of σ^B^ pulsing

In a recent study, Locke et. al. [13] demonstrated that a step-increase in energy stress results in pulsatile activation of σ^B^. The study also proposed a minimal mathematical model of the network which reproduced pulsing in σ^B^. However, this model included several assumptions inconsistent with experimentally observed details: (i) Phosphorylation and dephosphorylation reactions were assumed to follow Michaelis-Menten kinetics despite the fact that kinase (RsbW) and phosphatase concentrations are known to be comparable to substrate (RsbV) concentrations [18] so the approximation breaks down [19], (ii) σ^B^ and RsbV are represented as a single lumped variable rather than separate species and, (iii) partner-switching, and the formation and dissociation of various RsbW_2_ complexes were not included explicitly. Though this minimal model produces pulses resembling their experimental observations, it does not depict a biochemically accurate picture of the σ^B^ network. Consequently it cannot be used to uncover the design features that enable σ^B^ pulsing.

To understand the σ^B^ network response we built on our earlier study [15] to develop a detailed mathematical model that explicitly includes all known molecular interactions in the network. Note that we made one significant change to the model discussed in [15]. The model in [15] assumed that the synthesis rates for σ^B^ and its operon partners (RsbW and RsbV) follow the stoichiometry of their binding ratios (i.e. *RsbW_T_/B_T_* = 2 and *RsbW_T_/RsbV_T_* = 1; where *B_T_, RsbW_T_* and *RsbV_T_* represent total σ^B^, RsbW and RsbV concentrations respectively). However experimental measurements have shown that σ^B^, RsbW and RsbV are produced in non-stoichiometric ratios [18]. Accordingly, in contrast to our earlier study, we assumed σ^B^, RsbW and RsbV can be produced in non-stoichiometric ratios and studied how changes in relative synthesis rates of σ^B^ operon partners affect the response of the σ^B^ network to step-increases in energy stress phosphatase levels. We note that RsbX, a negative regulator of RsbTU phosphatase [20], is not included in our model. RsbX was excluded for simplicity since it is not essential for the pulsatile response of the σ^B^ network [14].

Simulations of this detailed model showed that different combinations of RsbW:σ^B^ and RsbV:σ^B^ relative synthesis rates lead to qualitatively different dynamical responses of the σ^B^ network. For operon partner synthesis ratios similar to those estimated in [18], our model responded to a step-up increase of the phosphatase with a pulsatile σ^B^ response (Fig. 1B) that resembled the experimentally observed behavior [13]. In contrast, when RsbW:σ^B^ and RsbV:σ^B^ relative synthesis rates follow the stoichiometry of their binding ratios pulsing is not observed and the σ^B^ activity monotonically increases over time (Figs. 1C). Pulsing also disappears when RsbW synthesis is high enough to neutralize both its binding partners (Figs.1D).

### Pulsing originates from emergent negative feedback in the network

To understand why the pulsatile response is only observed for certain operon partner synthesis rates, we investigated our mathematical model by decoupling the network’s transcriptional and post-translational responses. By varying the σ^B^ operon transcription rate, while keeping the relative synthesis rates of RsbW:σ^B^ and RsbV:σ^B^ fixed, we were able to calculate the post-translational response (Fig. 2A, blue curve) of the σ^B^ network: [σ^B^] = *F_P_* (*B_T_, P_T_*). This function describes how the free σ^B^ concentration ([σ^B^]) varies as a function of total σ^B^ (B_T_) and total phosphatase (P_T_) concentrations. In parallel, we calculated the transcriptional response (Fig. 2A, black curve) *B_T_ = F_T_* (*σ^B^*) which describes how changes in the free σ^B^ concentration affect total σ^B^ concentrations. In this analysis framework, the steady state of the complete closed loop network can be determined by simultaneously solving the post-translational and transcriptional equations, [σ^B^] = *F_P_* (*B_T_, P_T_*) and *B_T_ = F_T_* ([σ^B^]) at each phosphatase concentration *P_T_*. Graphing both functions provided the steady-state solution as their intersection point (Fig. 2A, red circle).

**Figure 2.**
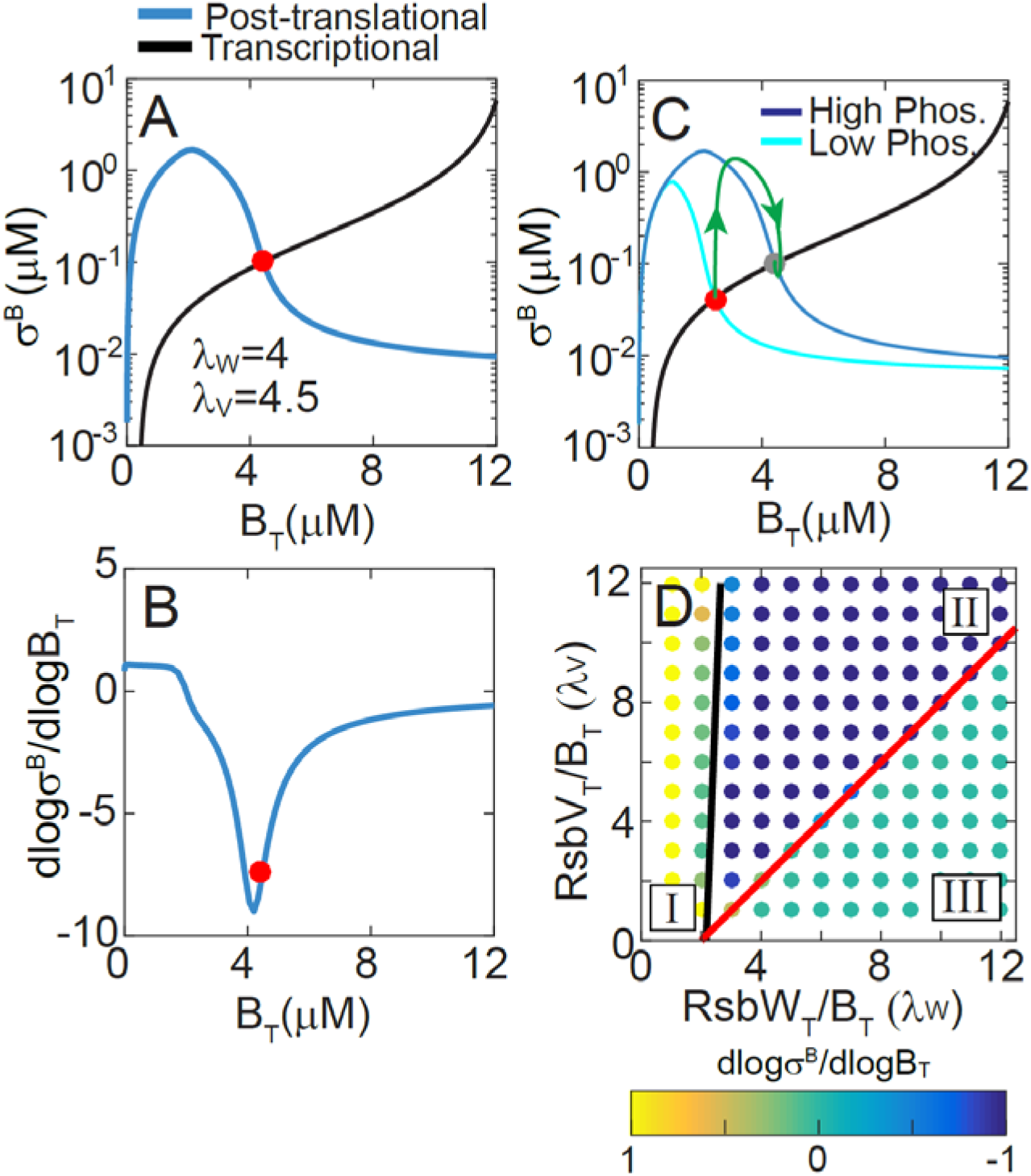
Negative feedback drives the pulsatile response of the σ^B^ network. **A.** Decoupled post-translational (blue curve) and transcriptional (black curve) responses of the σ^B^ network for *λ_W_* = *RsbW_T_/B_T_* = 4, *λ_V_ = RsbV_T_/B_T_* = 4.5. σ^B^ and Bt represent the concentrations of free and total σ^B^. Red circle marks the steady states of the full system. **B**. Sensitivity of the post-translational response (*LG_p_*) to changes in total σ^B^ concentration (operon production). **C**. Representation of the σ^B^ pulsatile trajectory in the σ^B^-B_T_ phase plane (green curve). Blue and cyan curves are the decoupled post-translational responses at high and low phosphatase concentrations. Black curve is the transcriptional response. **D**. (*λ_W_, λ_V_*) stoichiometry parameter space is divided into regions with positive (Region I), negative (Region II) and zero (Region III) post-translational sensitivity that respectively correspond to an effective positive, negative and no feedback in the σ^B^ network. Red and black lines represent the analytically calculated region boundaries *λ_W_* = 2 + *λ_V_* and *λ_W_* = 2(1 + *λ_V_k_deg_*/*k_k_*).

This decoupling approximation allows us to quantify the sign and strength of feedback in the full model. The effective sign of the feedback in the σ^B^ network is given by the sign of the product of the sensitivities of two response functions, i.e. sign((*∂F_T_*/*∂*[σ^B^])·(*∂F_P_*/*∂B_T_*)). Since σ-factors function as activators of transcription, *F_T_*(σ^B^) is a monotonically increasing function of σ^B^ (i.e. *∂F_T_*/*∂*[σ^B^]> 0). Consequently, the sign of the feedback in the σ^B^ network is given by the sign of the sensitivity of the post-translational response to B_T_ (i.e. *∂F_P_*/*∂B_T_*). In other words, if increase in the operon production leads to an increase in free σ^B^ then the feedback is positive, whereas if increase in the operon production leads to a decrease in free σ^B^ then the feedback is negative. Our results show that for the parameters chosen in Fig. 1B *F_P_* is a non-monotonic function of B_T_ (Fig. 2A, blue curve). At low B_T_, free σ^B^ increases as a function of B_T_ because RsbW is sequestered in the W_2_V_2_ complex. However at higher B_T_, the kinase flux dominates the phosphatase flux resulting in an increased RsbV~P and the freeing of RsbW_2_ from RsbV. Freed RsbW_2_ sequesters σ^B^ in the W_2_σ^B^ complex. Furthermore, in the total σ^B^ concentration range where ∂F_P_/∂B_T_ < 0 in Fig. 2B, the post-translational response is quite steep (Fig. 2A), i.e. small changes in B_T_ lead to significant decreases in free σ^B^. This ultrasensitivity can be quantified by calculating the slope in logarithmic space, i.e.

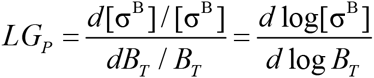

This dimensionless quantity characterizes the ratio of relative changes in σ^B^ and B_T_ at steady state (Fig. 2B). The sign of *LG_P_* defines the effective sign of the feedback loop and if the magnitude of |*LG_P_*| > 1 defines a super-linear (sigmoidal) response. For |*LG_P_*| ≫1 the response is ultrasensitive. For the σ^B^ network, in the region around the steady state *LG_P_<−1* indicating that the σ^B^ network operates in an ultrasensitive negative feedback regime. Two types of post-translational reactions that are known to produce ultrasensitivity play a role here (Fig. S1A,B): (1) Zero-order ultrasensitivity due to competition between RsbW kinase and RsbQP/RsbTU phosphatases for RsbV and (2) molecular titration due to sequestration of σ^B^ by RsbW. Therefore, near the steady state the σ^B^ network operates in an ultrasensitive negative feedback regime.

Notably, negative feedback is one of the few network motifs capable of producing adaption-like pulsatile responses [21]. Moreover, ultrasensitivity of the feedback ensures homeostatic behavior – making the steady state robust to variations of parameters [21]. This explains why in Fig. 1B a step-increase in the phosphatase concentration in our model leads to a σ^B^ pulse followed by return to nearly the same steady state. Plotting the trajectory of the σ^B^ pulse (green curve, Fig. 2C) on the ([*σ^B^*], *B_T_*) plane and over the post-translational and transcriptional responses (Fig. 2C) illustrates the mechanism driving this pulsatile response. Starting at the initial steady state (red circle), an increase in phosphatase shifts the ultrasensitive post-translational response (cyan to blue curve) so that free σ^B^ is rapidly released from the RsbW_2_-σ^B^ complex whereas total σ^B^ levels remain relatively unchanged. The increase in σ^B^ operon transcription eventually causes accumulation of total σ^B^ and the anti-σ-factor RsbW. This in turn forces the σ^B^ level to decrease, following the post-translational response curve, to the new steady state (gray circle) which has very little free σ^B^ thereby completing the σ^B^ pulse.

The same analysis can be applied for different values of relative synthesis rates, i. e. those that correspond to Fig. 1CD. As shown in Fig. S2 these parameter values do not produce an ultrasensitive non-monotonic post-translational response. Consequently they do not lead to the emergence of overall negative feedback explaining their non-pulsing dynamics. To determine if the presence or absence of negative feedback more generally explains the different dynamical responses in Fig. 1B-D, we sampled different combinations of relative synthesis rates (*RsbW_T_*/*B_T_* = *λ_W_* and *RsbV_T_*/*B_T_* = *λ*) and calculated the post-translational sensitivities. Our calculations showed that based on the sign of post-translational sensitivity (*LG_P_*) the relative synthesis parameter space can be divided into three regions (Fig. 2D). For (λ_W_, λ_V_) combinations in Region I the sensitivity is always positive. Increase in λ_W_ leads the system into an ultrasensitive negative regime (*LG_P_* < 0 and|*LG_p_*| ≫1) in Region II. A further increase in λ_W_ or a decrease in λ_V_ transitions the system into a non-responsive (*LG_P_* ~0) state in Region III. Dynamic simulations for sampled (λ_W_, λ_V_) combinations confirm that pulsatile responses to step-up in phosphatase concentration are restricted to Region II where the effective feedback is negative (Fig. S2).

To understand the boundaries between the three regions and how the level of the phosphatase affects the network, we developed a simplified analytical model that is based on the observation that RsbW and RsbV bind strongly to each other [18] (see Supporting Text for details). This approximation allowed us to determine the boundaries in Fig. 2D (black and red lines) and resulted in a clear biological interpretation of the three regions. In Region I the amount of RsbW, irrespective of phosphatase level, is insufficient to bind all of its partners and consequently some fraction of σ^B^ always remains free or unbound to RsbW. In contrast in Region II, the amount of phosphatase determines how much RsbV is in its inactive phosphorylated form RsbV~P and therefore whether the amount of RsbW is sufficient to bind all of its partners depends on the levels of RsbV~P. As a result, for this region, the ratio of kinase and phosphatase (P_T_) fluxes determines the post-translational response. Lastly, Region III is the opposite of Region I in that the amount of RsbW is more than sufficient to bind all of its partners, even when all RsbV is unphosphorylated. As a result, irrespective of phosphatase levels, very little σ^B^ is free and its level is nearly insensitive to changes in total σ^B^. Thus negative feedback and consequently pulsing are only possible in Region II where changes in phosphatase can shift the balance between the prevalent partner complexes.

The role of negative feedback in producing a pulsatile response also explains why pulsing does not occur in strains where σ^B^ operon is transcribed constitutively [13]. In this case, the σ^B^ network lacks the negative feedback necessary to produce a pulsatile response. A step-increase in phosphatase still leads to an increase in free σ^B^ due to the change in the post-translational response; however, this not followed by an increase in total σ^B^ levels (Fig. S2C). Consequently, an increase in phosphatase results in a monotonic increase in free σ^B^ rather than a pulse (Fig. S2F).

Further our decoupling method also sheds light on another experimental observation by Locke et. al. [13]: the dependence of σ^B^ pulse amplitude on the phosphatase level. Specifically, we found that σ^B^ pulse amplitude is a threshold-linear function of the phosphatase concentration (Fig. S3). Our decoupling method shows that this threshold-linear behavior arises because the σ^B^ network only operates in a negative feedback regime for phosphatase concentrations higher than a threshold. Below the phosphatase threshold, the post-translational response σ^B^ = *F_P_* (*B_T_, P_T_*)~0 and is insensitive to B_T_ (Fig. S3BC). Thus, the full system lacks the negative feedback and as a result σ^B^ does not pulse. Using our analytical approximation we found that this phosphatase threshold is proportional to the basal level of RsbW kinase synthesis rate and the ratio of the kinase and phosphatase catalytic rate constants (Fig. S3DE). Increase in the basal σ^B^ operon expression rate increases the phosphatase threshold. Further, an increase in the relative synthesis rate of RsbW (λ_W_=RsbW_T_/B_T_) makes the phosphatase threshold more sensitive to the σ^B^ operon expression rate, whereas a decrease in ratio of the kinase and phosphatase catalytic rate constants makes it less sensitive (Fig. S3DE). This shows that the phosphatase threshold represents the concentration at which the phosphatase is able to match the basal kinase flux.

Altogether these results show how the ultrasensitive negative feedback plays a critical role in determining many properties of the σ^B^ network pulsatile response and how the decoupling method can facilitate the identification of essential design features that enable the existence of this negative feedback.

### Under energy stress conditions σ^B^ network encodes phosphatase burst size into pulse amplitudes

In the preceding sections we have shown how the σ^B^ network responds to a step-increase in RsbQP or RsbTU phosphatases by producing a single pulse of activity. However, Locke et. al. [13] have shown that an increase in energy stress leads to a sustained response with a series of stochastic pulses in σ^B^ activity. This study further showed that this sustained pulsing response is driven by noisy fluctuations in level of energy-stress-sensing phosphatase RsbQP. While the mean level of RsbQP level is regulated transcriptionally by energy stress [8,13], its concentration in single cells can fluctuate due to the stochasticity of gene expression. To determine if our model could explain this response to stochastic fluctuations in RsbQP, we modified it to include fluctuations in the concentration of this phosphatase.

Based on previous theoretical [22,23] and experimental [24] studies we assume that fluctuating phosphatase level follows a gamma distribution which is described by two parameters - burst size (*b*, average number of molecules produced per burst) and burst frequency (a, number of bursts per cell cycle). The mean phosphatase in this case is the product of burst size and burst frequency 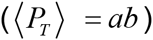. Thus, energy stress can increase mean phosphatase by changing burst size or burst frequency or both. In other words, stress conditions can increase phosphatase levels by either producing more phosphatase molecules per transcription-translation event or by making these events more frequent. While the results of [13] cannot exclude either mechanism, we can use our model to uncover which mechanisms is dominant.

First, we performed stochastic simulations in which mean phosphatase concentration was varied by changing burst size. These simulations reproduced all the experimentally-observed features of the σ^B^ pulsatile response. Specifically our results show that stochastic bursts in stress phosphatase levels lead to pulses of σ^B^ activity (Fig. 3A). Moreover, consistent with the experimental observations of [13], our model showed that the amplitude of σ^B^ pulses increases linearly with the stress phosphatase level (Fig. 3A,B). Finally, we found that stress-mediated increases in phosphatase concentration lead to an ultrasensitive (effective Hill coefficient ~5.6) increase in the frequency of σ^B^ pulsing (Fig. 3C) and an ultrasensitive (effective Hill coefficient ~2) increase in the level of σ^B^ target expression (Fig. 3D).

**Figure 3.**
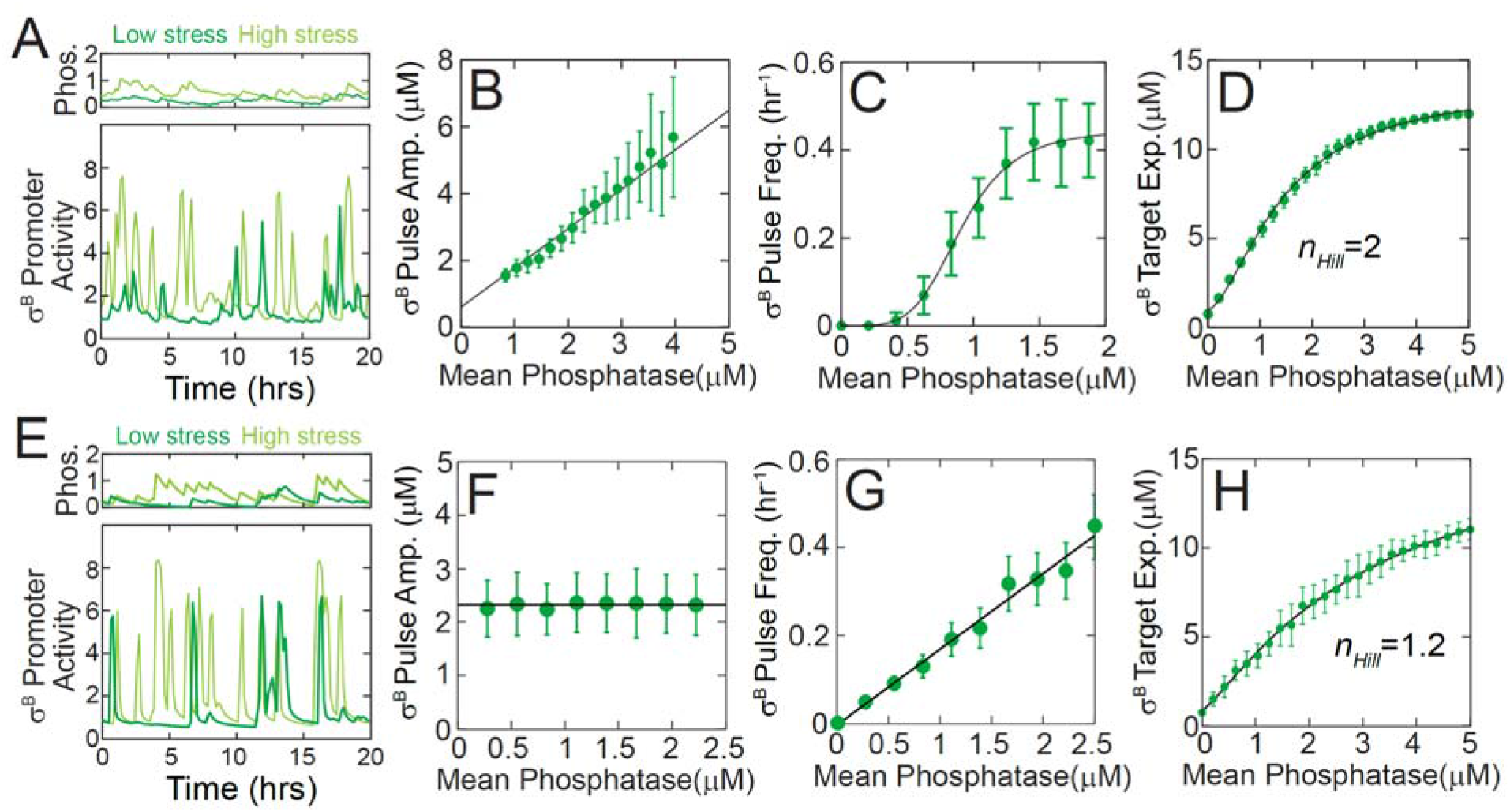
Pulsatile response of the σ^B^ network to stochastic phosphatase bursts during energy stress. Model simulations for σ^B^ network response where energy stress leads to an increase in stress-sensing phosphatase RsbQP burst size (A-D) or RsbQP burst frequency (E-H). **A,E**. Simulations show stochastic bursts in levels of RsbQP lead to pulses of σ^B^ target promoter activity. Light and dark green curves are sample trajectory from stochastic simulation at high and low stress respectively. Note that σ^B^ target promoter activity pulse amplitude increases significantly with increasing stress for burst size modulation (A) but not for burst frequency modulation (E). **B,F**. Mean σ^B^ pulse amplitude increases linearly as a function of mean phosphatase level for burst size modulation (B) but is insensitive to mean phosphatase level for burst frequency modulation (F). Green circles and errorbars show means and standard deviations calculated from stochastic simulations. Black line is a linear fit. **C,G**. With increasing mean phosphatase level, mean σ^B^ pulse frequency increases ultrasensitively for burst size modulation (C) and linearly for burst frequency modulation (G). Green circles and errorbars show means and standard deviations calculated from stochastic simulations. Black curves are a Hill-equation fit with *n_Hill_*=5.6 in (C) and a linear fit in (G) respectively. **D,H**. Mean σ^B^ target expression increases ultrasensitively as a function of mean phosphatase level for both burst size (D) and burst frequency (H) modulation. Green circles are the mean σ^B^ target expression calculated from stochastic simulations. Black curve is a Hill-equation fit with *n_Hill_* = 2 in (D) and in *n_Hill_* = 1.2 (H).

Next, we compared these results with stochastic simulations in which burst frequency was modulated (Fig. 3E-H). These simulations also led to an increase in σ^B^ pulsing (Fig. 3E) and a non-linear increase in the level of σ^B^ target expression as mean phosphatase level was increased with more frequent bursts (Fig. 3H). However, we found that σ^B^ pulse amplitude remains constant for burst frequency modulation (Fig. 3E,F) unlike the ~5-fold increase for burst-size modulation (Fig. 3B). Moreover, the frequency of σ^B^ pulses increase linearly with phosphatase level unlike the non-linear increase observed with burst-size-increase simulations (compare Figs. 3C and 3G). Notably the experimental observations reported in [13] show that σ^B^ pulse amplitude does increase (~3-fold) with an increase in energy stress thus suggesting that increase in phosphatase concentration at high stress is primarily the result of increase in burst size.

To further reinforce the role of mean burst-size modulation in controlling the σ^B^ pulsatile response we next examined the cumulative histograms of pulse amplitudes at different phosphatase concentrations. These histograms carry different signatures for burst-size or burst-frequency encoding. The distribution of pulse amplitudes is unchanged with increase in burst frequency (Fig. S4A) because σ^B^ pulse amplitude is determined by phosphatase burst size and not burst frequency. In contrast, if phosphatase levels are controlled by changing mean burst size then the distribution of pulse amplitudes changes accordingly. Consequently, the normalized cumulative histograms of pulse amplitudes overlap for burst-frequency encoding (Fig. S4A), but not for burst-size encoding (Fig. S4B). Applying this test to the data from [13], we found that the normalized cumulative pulse amplitudes histograms do not overlap (Fig. S4C). These results predict that stress affects the σ^B^ network via burst-size modulation of phosphatase production which is then encoded into σ^B^ pulse amplitudes. While the molecular mechanism that introduces energy stress to the network is still not fully understood, our prediction places an important constraint on it.

### σ^B^ network encodes rate of environmental stress increase into pulse amplitudes

Our model can also be used to study the response of σ^B^ network to environmental stress. Unlike the energy stress phosphatase, the environmental stress phosphatase RsbU is regulated post-translationally by binding of RsbT [25–27]. RsbT is trapped by its negative regulators under unstressed conditions but is released upon stress. Consequently, the concentration of RsbTU complex is tightly controlled at the post-translational level and is therefore expected to be relatively insensitive to gene expression fluctuations but sensitive to the level of environmental stress. As a result, step-up increases in environmental stress agents like ethanol produce rapid increases in RsbTU and result in only a single pulse of σ^B^ activity [14]. However it has been shown that for gradual increases in stress, σ^B^ pulse amplitude depends on the rate of stress increase [14]. To explain this response, we modeled gradual stress with ramped increase in RsbTU complex concentration (Fig. 4A). Our simulations showed that the detailed model of σ^B^ network is indeed able to capture the effect of rate of stress increase on σ^B^ pulse amplitudes. Specifically for a fixed increase in RsbTU complex, the pulse amplitude decreases non-linearly as a function of the duration of phosphatase ramp (Fig. 4B, E).

**Figure 4.**
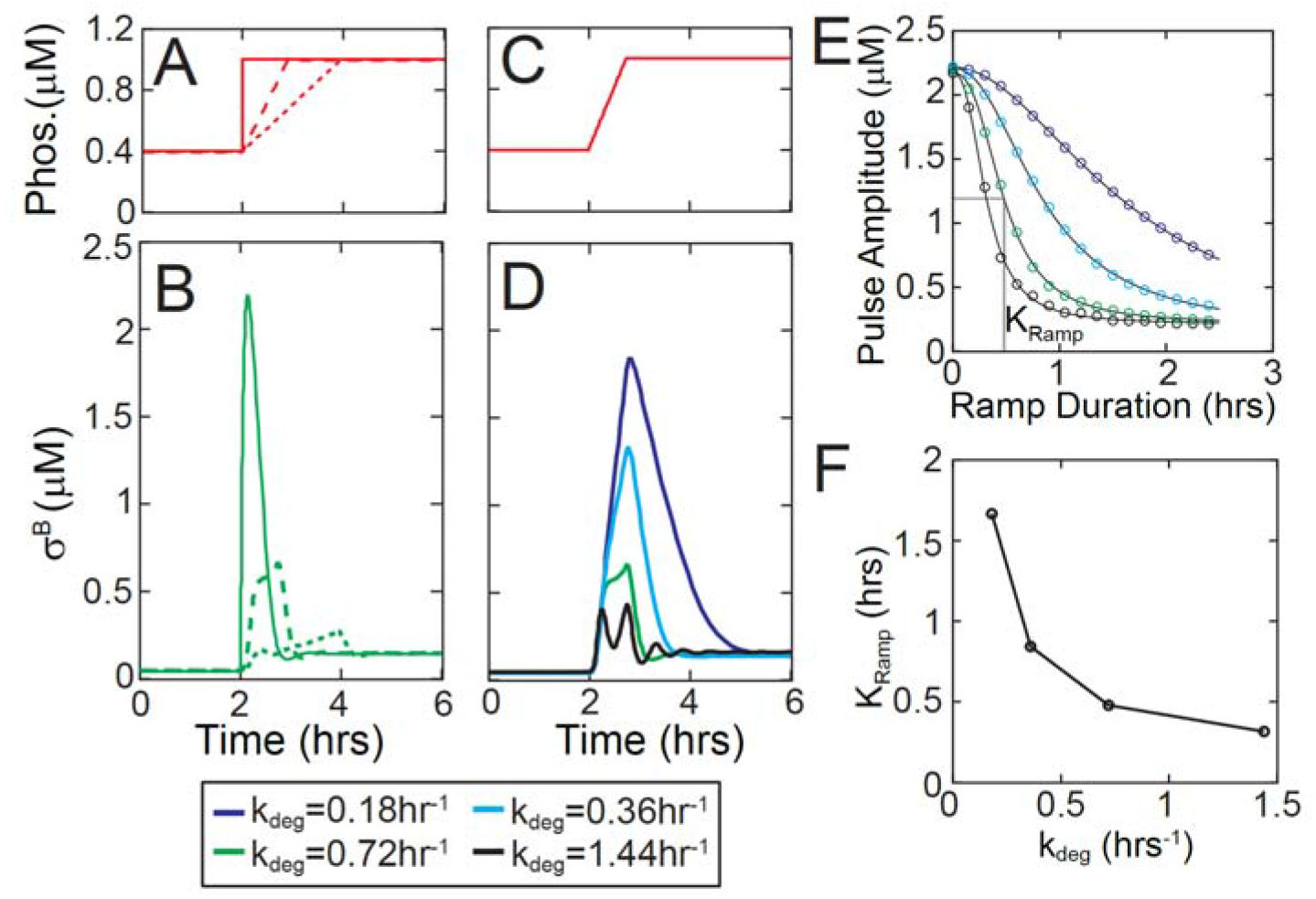
Rate sensitivity of the σ^B^ pulsatile response to environmental stress. **A.** Ramped increases in RsbTU complex concentration were used as model inputs to simulate different rates of stress increase in σ^B^ network. **B**. σ^B^ pulse amplitudes in the wildtype model (k_deg_ = 0.72 hr^−1^ is the degradation rate of σ^B^ operon proteins) resulting from the ramped increases in phosphatase concentration shown in (A). **C,D**. σ^B^ pulse amplitudes resulting from the ramped increase in phosphatase concentration shown in (C) for various degradation/dilution rates (D). **E**. Non-linear dependence σ^B^ pulse amplitude on phosphatase ramp duration for various degradation/dilution rates. Circles and solid curves represent simulation results and Hill-equation fits respectively. Colors represent different k_deg_ values as in (D). **F**. K_ramp_, the half-maximal constant of the non-linear dependence of amplitude on ramp duration, as a function of k_deg_.

We hypothesized that this ramp rate encoding is the result of the timescale separation between the fast post-translational and the slow transcriptional responses of the σ^B^ network. During the pulsed σ^B^ activation, post-translational response is rate-limited by the phosphatase ramp. In contrast, the transcriptional response is slow and its rate is set by the degradation rate of σ^B^ operon proteins. Following a step-increase in phosphatase, the fast post-translational response ensures that σ^B^ reaches its post-translational steady state before the slow increase in RsbW sequesters σ^B^ and turns off the pulse (Fig. 4AB). However, for a ramped increase in phosphatase the post-translational increase in σ^B^ is limited by the rate of phosphatase ramp. This allows RsbW to catch up and terminate the σ^B^ pulse earlier, thereby decreasing the pulse amplitude. To test this, we varied the degradation rate of σ^B^ operon proteins and proportionally changed the operon transcription rate to ensure that the total concentrations of σ^B^, RsbW and RsbV are kept fixed. We found that indeed pulse amplitude decreases with increase in degradation/dilution rate (Fig. 4CD). Our simulations showed that K_ramp_, the half-maximal constant for the dependence of pulse amplitude on ramp duration, was indeed sensitive to the degradation rate (Fig. 4EF). This suggests that the timescale separation between the post-translational and transcriptional responses is the basis of ramp rate encoding into pulse amplitude.

### The design of the σ^B^ network enables it to compete with σ^A^ for RNA polymerase

The results thus far indicate that σ^B^ network functions in the effectively negative feedback regime where increase in the operon expression decreases σ^B^ activity. Negative feedback loops have been shown to increase the robustness of the system to perturbations. We therefore decided to investigate how the σ^B^ network design affects its performance when it faces competition for RNA polymerase from other σ-factors, e.g. from the housekeeping σ-factor σ^A^ [16,28,29]. Since σ^A^ has a much higher affinity for RNA polymerase [17], a small increase in σ^A^ can dramatically increase the amount of σ^B^ necessary to activate the transcription of the σ^B^ regulon. Thus, changes in σ^A^ can alter the input-output relationship of a stress-response σ-factor like σ^B^ (Fig. S5AB) and thereby adversely affect the survival of cells under stress.

To understand how the σ^B^ network handles competition for RNA polymerase, we expanded our model to explicitly include σ^A^, RNA polymerase (RNAPol) and its complexes with both σ-factors. The presence of σ^A^ will affect transcriptional activity of σ^B^ but not post-translational interactions between σ^B^ operon partners (Fig. 5A, left panel). Therefore, post-translational response σ^B^ = *F_P_* (*B_T_, P_T_*) is not affected by σ^A^. In contrast, in the transcription response, an increase in σ^A^ decreased the ‘effective affinity’ of σ^B^ for RNApol and consequently higher levels of free σ^B^ are necessary to achieve the same production rate for σ^B^ target genes.

**Figure 5.**
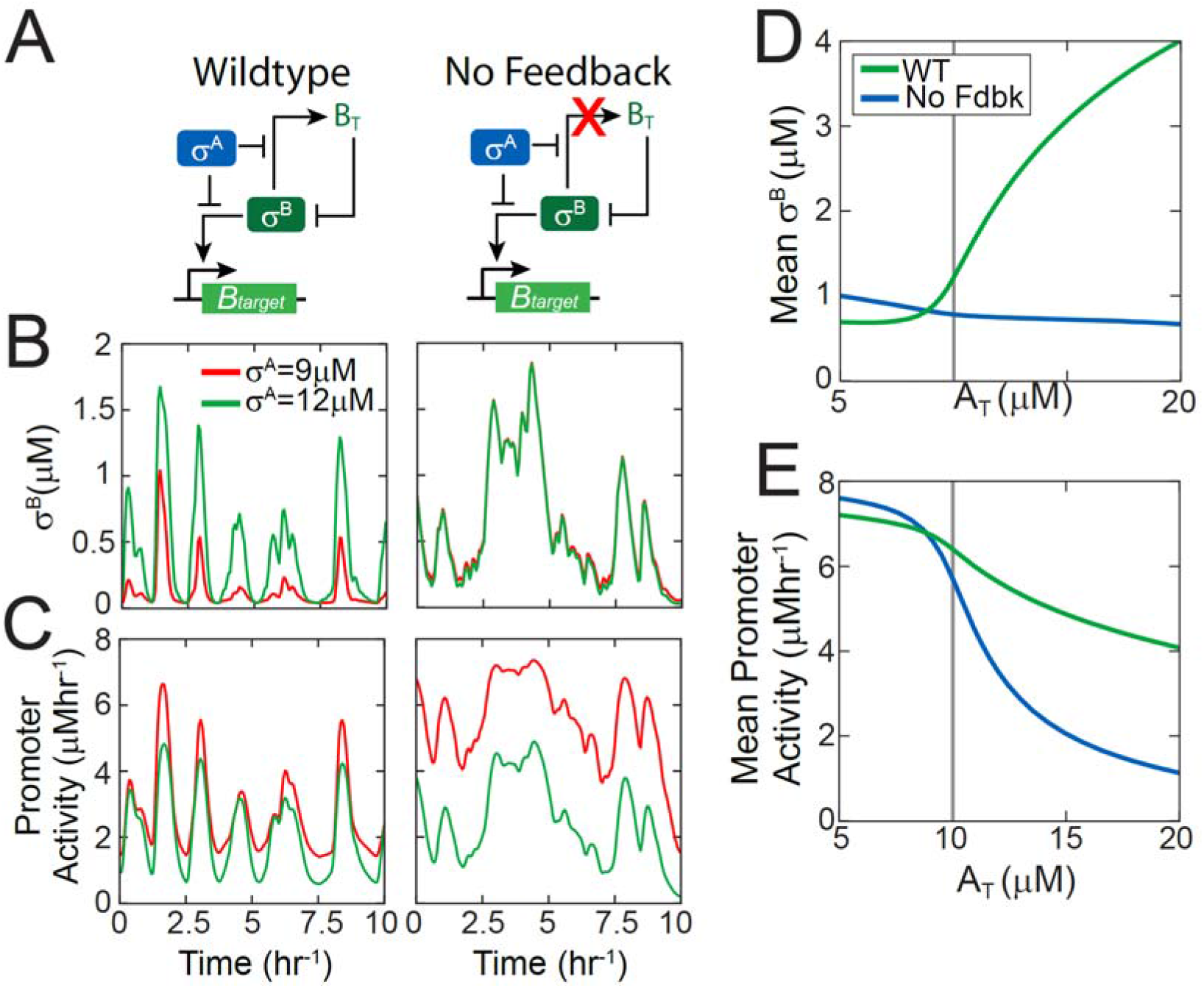
Negative feedback insulates the σ^B^ response from competition with houskeeping σ-factor σ^A^. **A.** Simplified network diagrams of stress σ-factor σ^B^ competing with housekeeping σ-factor σ^A^ for RNA polymerase. In all cases, a σ^B^ phosphatase controls the stress-signal driven activation of σ^B^. **(B,C)**. Trajectories of free σ^B^ (B) and σ^B^ target promoter activity (C) in response to stochastic phosphatase input for both networks at two different levels of σ^A^ (σ^A^ = 9μM – low competition-regime and σ^A^ = 12μM – high-competition regime for RNA polymerase). **D-E**. Mean free σ^B^ concentration (D) and mean σ^B^ target promoter activity (E) as a function of total σ^A^ concentration (A_T_) for both networks in (A) at fixed mean phosphatase (mean P_T_ = 0.5 μM). Gray vertical line shows the total RNA polymerase level which was fixed at 10 μM.

Using our model, we examined how changes in σ^A^ level affect the network response to energy stress signal, i.e. under stochastically fluctuating RsbQP phosphatase levels. Our simulations showed that phosphatase bursts lead to pulses of free σ^B^ and pulsatile transcription of σ^B^-controlled promoters (Fig. 5BC) as the presence of σ^A^ does not affect the effective feedback sign. Notably our results also showed that the amplitudes of σ^B^ target promoter pulses are hardly affected by a ~30% increase in σ^A^ (Fig. 5C, left panel). This surprising insensitivity of the phosphatase-σ^B^ target dose-response to RNApol competition is the result of the ultrasensitive negative feedback between free σ^B^ and total σ^B^. Due to the ultrasensitivity of this feedback, a small decrease in total σ^B^ levels resulting from the increase in σ^A^ causes a large increase in σ^B^ pulse amplitude (Fig. 5B left panel, 5D green line). This increased amplitude compensates for the increased competition for RNApol and insulates the network from perturbations (Fig. 5DE, green curves).

To further illustrate the importance of the negative feedback in insulating the network, we compared the response of the wildtype network to an *“in silico”* mutant network wherein the σ^B^ operon is constitutive rather than σ^B^ dependent (Fig. 5A). Consequently this network lacks any feedback between free σ^B^ and total σ^B^. Our simulations (Fig. 5B, right panel) show that the free σ^B^ concentration of the no-feedback-network does not show adaptive pulsing and therefore σ^B^ concentration fluctuates along with the phosphatase levels. Increase in σ^A^ did not affect this response. This is expected since in the absence of feedback σ^A^ only affects the expression of σ^B^ targets in this network (Fig. 5A, right panel). Without an increase in free σ^B^ (Fig. 5D), the increased competition for RNApol at higher σ^A^ reduced the σ^B^ target promoter activity (Fig. 5CE). Similarly a positive feedback network design is also incapable of increasing free σ^B^ in response to an increase in σ^A^ (Fig. S5CDE). Thus fluctuations in σ^A^ can interfere with the σ^B^ stress-response of these alternative network designs. In contrast, the wildtype σ^B^ network with its ultrasensitive negative feedback design can compensate for competition effects (Fig. 5DE).

### Negative feedback designs of stress-response σ-factor networks minimizes interference

The emergent negative feedback design of the network discussed here is not unique to σ^B^. Transcription of many alternative σ-factors in *B. subtilis* as well in other bacteria is often positively auto-regulated but sigma-factor operons often include post-translational negative regulators [3,12,30–33]. For example σ^W^, a σ-factor in *B. subtilis* that controls the response to alkaline shock [34] is co-transcribed with its anti-σ-factor RsiW. In the absence of stress, RsiW sequesters σ^W^ in an inactive complex. σ^W^ is activated by stress signals which trigger the cleavage and degradation of RsiW thereby releasing and activating σ^W^ target expression [35]. Although it is unknown whether the σ^W^ network functions in a negative feedback regime similar to σ^B^ or if it pulses, it is possible for this network to exhibit these design properties. If RsiW is expressed in stoichiometric excess of its binding partner σ^W^ from the σ^W^-regulated operon which they share [36], then similar to the σ^B^ network, σ^W^ would operate in a negative feedback regime.

To determine if negative feedback control offers any advantages when multiple stress σ-factors are active, we built a new model that includes three σ-factors: σ^B^, σ^W^ and σ^A^. Anti-σ-factors RsbW (RsiW) and other details of post-translational regulation were excluded for simplicity. Instead the regulation of free σ^B^ and σ^W^ was modeled with simplified identical versions of the negative feedback design of the σ^B^ network (Fig. S5A). Under this simplification, free σ^B^ and free σ^W^ are non-monotonic functions of their respective total concentrations, *B_T_* and *W_T_*. These non-monotonic functions are qualitatively similar to the post-translational response function shown in Fig. 2B and depend on a signaling proteins P_B_ (for σ^B^) and P_W_ (for σ^W^). Following the previous section, this model explicitly includes σ^A^, RNApol and its complexes with σ-factors. As a result, transcriptional activity of both σ^B^ and σ^W^ depend on σ^A^ and RNApol concentrations (see Supplementary Text). Concentrations of RNApol and σ^A^ were chosen to ensure that amount of RNApol is insufficient to bind to all σ-factors at the same time. All other parameters of the simplified model were chosen to approximately match the full σ^B^ network model and ensure that both σ^B^ and σ^W^ operate in the negative feedback regime. Consequently for the chosen parameters this simplified model acts like our detailed model and responds to step increases in the stress signaling protein P_B_ (or P_W_) by producing a pulse of σ^B^ (or σ^W^) activity (Fig. S5CD). To enable a comparison of the competition between σ-factors for different types of feedback we hereafter focus on only steady state response, however our conclusions are also valid for the averaged pulsatile dynamical responses that could be characteristic of the negative feedback σ-factor networks.

We used this simple model to study the response when cells are simultaneously exposed to multiple stresses creating competition for RNApol. For these simulations we fixed σ^A^ levels and studied how activation signals for one alternative σ-factor affects the activity of another. As before (Fig. S5AB), increased availability of one stress σ-factor leads to a competition for RNA polymerase and as a result reduces the activity of another stress σ-factor (Fig. S6EF). However, when negative feedback loops are present, surprisingly, increasing the stress signal for one σ-factor did not lead to any significant change in the activity of another σ-factor. For example, increasing stress signaling protein P_B_ while keeping P_W_ fixed leads to an increase in free σ^B^ but also results in a small increase in free σ^W^ (Fig. 6C). This response can be explained by the ultrasensitive negative feedback loops controlling the two stress σ-factors. An increase in free σ^B^ by stress signaling protein P_B_ leads to increased competition for RNApol resulting in a decrease in the production of RsbW. But since σ^W^ is regulated by a negative feedback, a decrease in total RsbW concentration actually frees up more σ^W^ thereby insulating σ^W^ target activity from the effects of RNApol (Fig. 6E). Similarly the dynamic response of the stress σ-factors is also insulated from competition and an increase in fixed P_W_ levels increases the pulse amplitude of σ^B^ in response to step changes in stress signaling protein P_B_ (Fig. S6A-D). This compensation of changes in RNA polymerase availability comes about because both σ^B^ and σ^W^ are regulated by ultrasensitive negative feedbacks in our model. As a result of this negative feedback, both σ-factor networks function as homeostatic modules. Homeostatic resistance to changes in signals is an intrinsic property of ultrasensitive negative feedback motifs.

**Figure 6.**
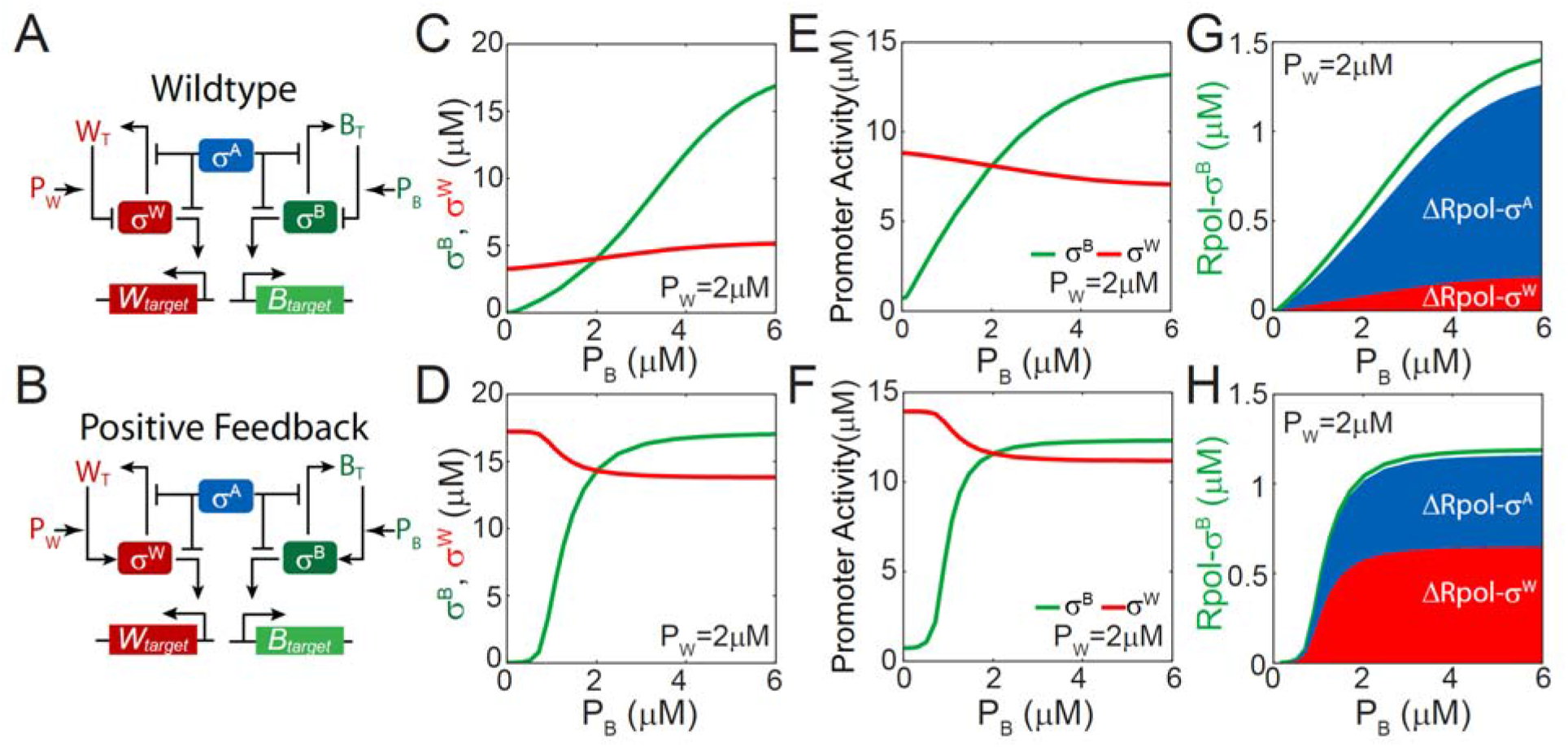
Negative feedback minimizes competition between stress σ factors for RNA polymerase. A,B. Simplified network diagrams of stress σ-factors σ^B^ and σ^W^ and housekeeping σ-factor σ^A^ competing with each other for RNA polymerase. σ^B^ and σ^W^ activities are regulated by negative and positive feedbacks in (A) and (B) respectively. In both cases, signaling proteins P_B_ and P_W_ control the stress-signal driven activation of σ^B^ and σ^W^ respectively. **C, D**. Dependence of free σ^B^ and σ^W^ levels on P_B_ at fixed P_W_ (= 2μM). In the wildtype negative feedback system (C), increase in σ^B^ phosphatase leads to an increase in both free σ^B^ (green curve) and free σ^W^ (red curve). In the positive feedback system (D), increase in σ^B^ phosphatase leads to an increase in free σ^B^ (green curve) and a decrease in free σ^W^ (red curve). **E, F.** σ^B^ and σ^W^ target promoter activities as a function of P_B_ at fixed P_W_ in the wildtype negative feedback system (E), and the positive feedback system (F). **G, H**. RNA polymerase bound σ^B^ (Rpol-σ^B^) as a function of P_B_ at fixed P_W_ in the wildtype negative feedback system (G) and the positive feedback system (H). Increase in σ^B^ phosphatase (P_B_) leads to an increase in Rpol-σ^B^ (green curve) and corresponding decreases ΔRpol-σ^W^ in Rpol-σ^W^ (red area) and ΔRpol-σ^A^ in Rpol-σ^A^ (blue area).

Thus the two stress σ-factors are able to function simultaneously despite the scarcity of RNApol. The mechanism minimizing competition between stress σ-factors becomes clearer when we track the changes in σ–RNApol complexes as a function of the stress signaling protein P_B_. As P_B_ increases, more free-σ^B^ becomes available and binds to RNApol (Fig. 6G). However this RNApol must be accounted for by the RNApol lost by the other operating σ^W^ and σ^A^ factors. Comparing the contributions of each σ-factor shows that despite the fact that σ^A^ has a much higher affinity for RNApol, most of the RNApol in the σ^B^-RNApol complex is drawn from the σ^A^-RNApol pool rather than σ^W^-RNApol pool (Fig. 6G). Thus the negative feedback design allows stress σ-factors to minimize their competition with each other at the expense of the housekeeping factor σ^A^.

The role of the negative feedback in producing this response becomes clear when we compare the response of an *“in silico”* mutant network with positive feedback loops between σ^B^ and B_T_ and σ^W^ and W_T_. These positive feedback loops are expected to display no homeostatic properties and as a result, in this network activation of σ^B^ should significantly decrease σ^W^ activity. Indeed, our simulation for the positive feedback network (Fig. 6D) demonstrates that with increase in stress signaling protein P_B_ and the resulting increase in free σ^B^, the free σ^W^ concentration decreases. As a result of the increased competition for RNApol and the decreased free σ^W^, σ^W^ target promoter activity in this network decreases as a function of P_B_ (Fig. 6F). Moreover comparing changes in σ-RNApol complexes as a function of stress signaling protein P_B_ we find that most of the RNApol in the σ^B^-RNApol complex is drawn from the σ^W^-RNApol pool rather than σ^A^-RNApol pool (Fig. 6H). Thus the negative feedback designs are essential for stress σ-factors not only to tolerate competition from σ^A^, but also to avoid competing with each other when the cell is simultaneously exposed to multiple types of stresses.

## Discussion

Taken together, our results show how the design of the σ^B^ network includes an implicit ultrasensitive negative feedback that plays multiple functional roles. This design enables pulsatile activation of σ^B^ in response to energy stress and rate-sensitivity to increases in environmental stress. Moreover, our model predicts that the same design feature allows the network to effectively compete with house-keeping and other alternative σ-factors for RNA polymerase core.

Prompted by recent observations of the highly dynamic pulsatile response of the σ^B^ network [13,14], we have developed a mathematical model that reproduces all reported features of the response including pulsatile activation in response to stress. Our model avoids making ad hoc simplifications and instead captures all the known molecular details of the network. By decoupling the post-translational and transcriptional responses in our model we were able to derive a simplified view of the network that illustrates how the pulsatile response is mechanistically based on the ultrasensitive negative feedback in the network. Using this method we identified the relative stoichiometry of σ^B^, RsbW and RsbV synthesis rates as the most critical design property, which by controlling the post-translational response determines the sign of the feedback in the network as well as all qualitative features of the network response. This highlights how ignoring non-transcriptional interactions and focusing on transcriptional regulatory interactions alone can be misleading when trying to identify or characterize network motifs. Notably, recent analyses of networks like bacterial two-component systems [37] and the sporulation phosphorelay [38] have similarly shown how the effective sign of feedback in these networks depends critically on their post-translational interactions.

The decoupling of the post-translational and transcriptional response greatly facilitated the identification of critical design features despite the complexity of the network. This separation greatly reduces the dimensionality of the dynamical system by enabling an independent input–output analysis for the two modules. Similar methods have also been applied to deduce core functional properties in other bacterial networks comprising two-component systems and alternative σ-factors [39–41]. Interestingly our analysis revealed that the post-translational and transcriptional module structures of the σ^B^ network and the phosphorelay controlling *B. subtilis* sporulation are remarkably similar [38]. Despite the differences in molecular details, in both networks increase in total transcription factor levels produces a non-monotonic response in the active transcription factor. Combining this response with the transcriptional feedback produces an ultrasensitive negative feedback in both networks. The relevance of these similarities is evidenced by the fact that both networks produce dynamically similar pulsatile responses even though they are activated by entirely different stimuli.

We further showed that energy stress can control σ^B^ pulses frequency by modulating the size of stochastic bursts of energy stress phosphatase. This result raises the question whether pulsatile σ^B^ response can achieve proportional expression of downstream genes, as was previously suggested [13,42]. This proportional control requires the distribution of pulse amplitudes to remain fixed even as stress levels increase. However under the burst-size encoding strategy, pulse amplitude distributions change as stress levels increase thereby negating the efficacy of a pulsed response in producing proportional expression of downstream genes. The functional significance of pulsatile response may instead lie in its ability to encode the rate of environmental stress increase. Our model showed that this rate encoding follows from the timescale separation between the fast post-translational and the slow transcriptional responses in the network. As a result cells are able to encode the rate of stress increase into σ^B^ pulses. This rate responsiveness is only possible with adaptive pulsatile responses and thus may explain the need for σ^B^ pulsing to control the general stress response.

We also used our model to understand the response when placed in the larger context of other σ-factor networks and competition for RNA polymerase. Our results show how the network design is uniquely suited to insulating its response from RNA polymerase competition from the housekeeping σ-factor. Finally we demonstrate how ultrasensitive negative feedback, a ubiquitous feature of stress σ-factor regulation enables different stress σ-factors to operate simultaneously without inhibiting each other. These results are relevant not only for understanding the stress response of bacteria but also increasingly for the design of synthetic circuits. The movement towards the construction of larger genetic circuits has produced numerous recent designs that include multiple independent modules that rely on shared resources or actuators to function [43–45]. Our results highlight how competition between modules for shared resources can significantly affect the performance of these synthetic circuits. Further, inspired by the design of naturally occurring stress σ-factor network we provide new design rules that can improve the performance and robustness of the synthetic networks.

## Acknowledgments

The research was supported by NIH grants GM 096189 to OAI. The authors are grateful to James Locke for sharing raw data from Ref. [13] and Chet Price for feedback on the manuscript.

## Methods

### Mathematical model of the σ^B^ network

The details of all biochemical reactions in the model and the corresponding differential equations are described in the Supplementary Text.

### Mathematical model of σ^B^ stress-response network

Our mathematical model of σ^B^ network is based on a previous model proposed in [15]. This ODE-based model explicitly includes all known molecular species, post-translational reactions and the transcriptional regulation of the σ^B^ operon by σ^B^. Below we formulate the set of reactions and associated differential equations.

### Model reactions

The events shown in Figure 1A can be described by the following set of biochemical reactions:

- Dimerization of anti-σ-factor RsbW

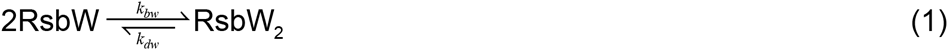
- Reversible binding of the anti-anti-σ-factor RsbV to anti-σ-factor dimer RsbW_2_ to form the complexes RsbW_2_-RsbV and RsbW_2_-RsbV_2_

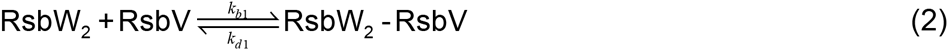

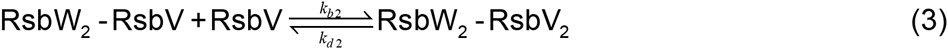
- Phosphorylation of the anti-anti-σ-factor RsbV by RsbW_2_

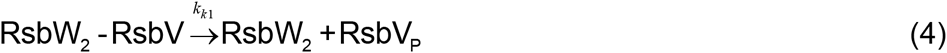

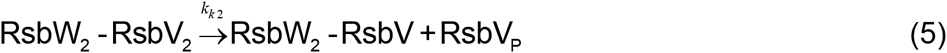
- Reversible binding of σ^B^ to RsbW_2_ to form the complex RsbW_2_-σ^B^

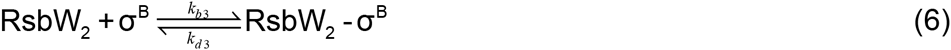
- Reversible displacement of σ^B^ by RsbV in the complex RsbW_2_

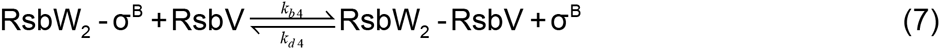
- Dephosphorylation of phosphorylated anti-anti-σ-factor RsbV~P

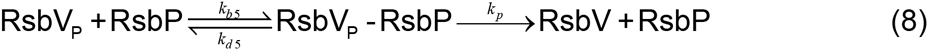
- Protein degradation/dilution due to cell growth

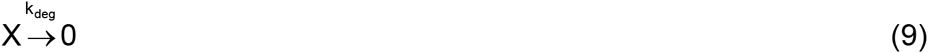

where X is any protein or protein complex in the σ^B^ network. For simplicity equal rates of degradation for all proteins and their complexes are assumed.
- Production of σ^B^, RsbW and RsbV

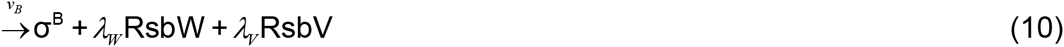

σ^B^, RsbW and RsbV were assumed to be synthesized proportionally as all three are part of the same operon. *λ_W_* and *λ_V_* are the proportionality constants of the relative synthesis rats of operon genes. Synthesis was modeled as a hyperbolically increasing function of σ^B^ concentration, [*σ^B^*], due autoregulation:
.

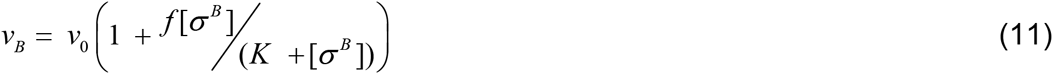

Here *v*_0_ is the basal synthesis rate, *f* is the fold change in protein synthesis due to positive autoregulation and *K* is the equilibrium dissociation constant for the binding of σ^B^ to the promoter DNA.

The stress signals were assumed to control the concentrations of stress phosphatases RsbTU and RsbQP. For RsbQP, energy stress was assumed to regulate the transcription rate of the phosphatase and the phosphatase concentration was assumed to be subject to stochastic fluctuations resulting from gene expression noise. In contrast, RsbTU concentration is regulated by environmental stress post-translationally, consequently RsbTU concentration was assumed to be stress-dependent but not subject to stochastic fluctuations.

### Model equations

We assume mass-action kinetics for all the above reactions (equations 1–10) to obtain the following set of equations that describe network dynamics:

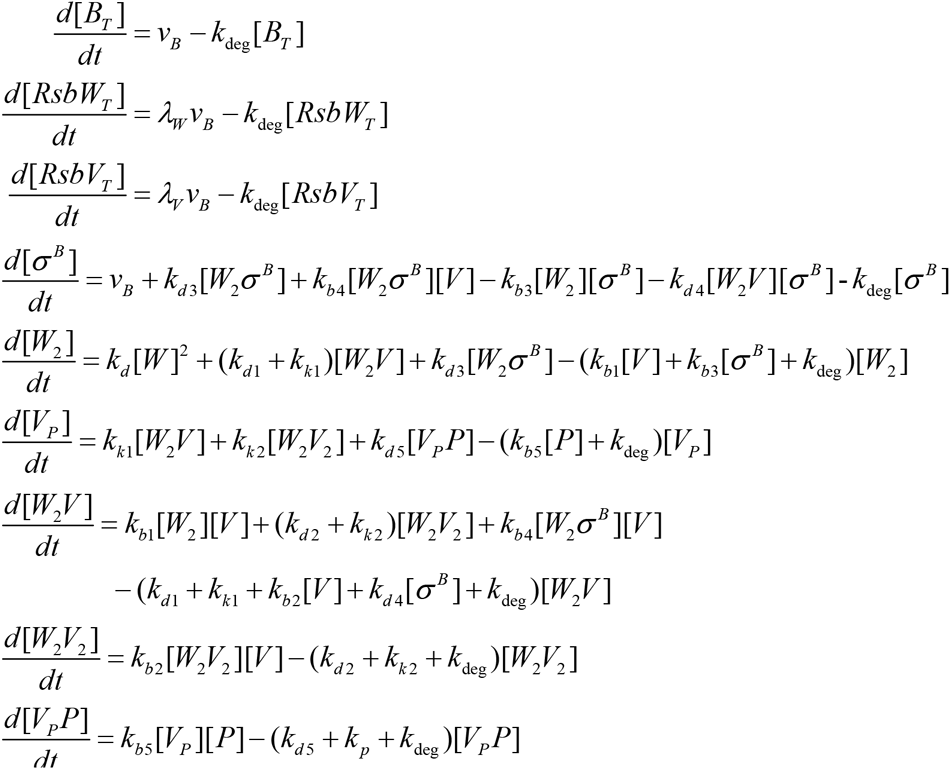

Here [*σ^B^*] is the concentration of free σ^B^; [*W_2_*] is the concentrations of dimeric RsbW; [*V*] and [*V_P_*] are the concentrations of unphosphorylated and phosphorylated RsbV; [*W*_2_*σ^B^*], [*W*_2_*V*], [*W*_2_*V*_2_] and [*V_P_P*] are the concentrations of the corresponding protein complexes. [*B_T_*], [*RsbW_T_*], [*RsbV_T_*] and [*P_T_*] are the concentrations of total σ^B^, RsbW, RsbV and phosphatase:

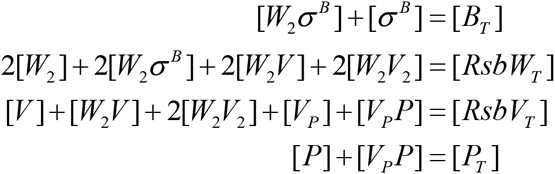

All model parameters are summarized in Table 1.

To study the effects of competition for RNA polymerase, the σ^B^ network model was expanded to include reactions for σ^A^, RNA polymerase (RNApol) and σ–RNApol binding (see Supplementary Text). To investigate the competition between σ^B^, σ^W^ and σ^A^, we used a phenomenological non-monotonic function to model the post-translational regulation of stress σ-factors (σ^B^ and σ^W^; see Supplementary Text for details).

### Calculation of steady state post-translational and transcriptional responses

The decoupled transcriptional and post-translational responses of the network at steady state were calculated using the MATLAB bifurcation package MATCONT. The post-translational response [*σ^B^*] = *F_P_* (*B_T_,P_T_*), was calculated by varying the rate of operon transcription while keeping the component synthesis rates (λ_W_, λ_V_) and the total phosphatase concentration (P_T_) fixed. Similarly, the transcriptional response B_T_ = F_T_(σ^B^), was calculated by varying the free σ^B^ concentration as an independent variable to calculate the total concentrations of σ^B^, RsbW and RsbV.

## Simulations

The parameter values for reversible binding and phosphorylation reactions were taken from [15] or were analysis driven to obtain pulsing in σ^B^. All the parameters used in the model are summarized in Table S1. In the deterministic set-up (Figs. 1, 2, 4, 6, S1, S2, S3 and S6) the system of differential equations was solved using standard *ode15s* solver in MATLAB. For stochastic simulations in Figs. 3, 5 and S5, the time-varying total phosphatase level P_T_ (=P + V_P_P) was pre-computed using a gamma distributed Ornstein-Uhlenbeck process as in [13]. This gamma distributed Ornstein-Uhlenbeck process permits independent modulation of mean burst size (*b*) and frequency (a) [46]. For each phosphatase level, 50 simulations were performed each lasting 10 hours. Pulses were detected by examining local maxima and minima of the simulated trajectories, and subsequently this information was used to compute statistics for pulse amplitude and frequency.

For the simulations of the effect of competition for RNA polymerase (Figs. 5 and S5), the total housekeeping σ-factor concentration was varied between 5 and 15 μM. In these simulations we used (λ_W_ = 4, λ_V_ = 4.5) and (λ_W_ = 2, λ_V_ = 2) to simulate the wildtype (negative feedback) and positive feedback networks respectively. For the simulations of the no feedback network we used (λ_W_ = 4, λ_V_ = 4.5) and f = 0 and v_0_ = 8.64 μMhr^−1^ to model the σ^B^–independent constitutive production of operon components.

For the simulations of the competition between σ^B^, σ^W^ and σ^A^ (Figs. 6 and S6), the total housekeeping σ-factor concentration was kept fixed at 12 μM. We used (nb = 7, mb = 5) and (nb = 0, mb = 3) to simulate the wildtype (negative feedback) and positive feedback networks respectively. K_B_ and K_W_ were fixed at 5μM for simulations of both networks.

**Table 1.** List of parameters values used in the model for σ^B^ network

| Parameter | Value | References |
| --- | --- | --- |
| $k_{bw}$ | $72 \mu\text{M}^{-1}\text{hr}^{-1}$ | [15,18] |
| $k_{dw}$ | $18 \text{ hr}^{-1}$ | [15,18] |
| $k_{b1}, k_{b2}, k_{b3}, k_{b5}, k_{bb}, k_{ba}, k_{bpb}$ | $144 \mu\text{M}^{-1}\text{hr}^{-1}$ | [15] |
| $k_{d1}, k_{d2}, k_{d3}, k_{d5}$ | $18 \text{ hr}^{-1}$ | Assuming binding affinity of $\sigma^B$ operon partner complexes $\sim 8\text{nM}$ [18] |
| $k_{b4}$ | $72 \mu\text{M}^{-1}\text{hr}^{-1}$ | [15] |
| $k_{d4}$ | $72 \mu\text{M}^{-1}\text{hr}^{-1}$ | [15] |
| $k_{db}$ | $172.8 \text{ hr}^{-1}$ | RNApol- $\sigma^B$ dissociation constant assuming a binding affinity of $1.2\mu\text{M}$ [16] |
| $k_{da}$ | $2.88 \text{ hr}^{-1}$ | RNApol- $\sigma^A$ dissociation constant assuming a binding affinity of $0.02\mu\text{M}$ [16] |
| $k_{dpb}$ | $14.4 \text{ hr}^{-1}$ | RNApol- $\sigma^B$-$p_B$ dissociation constant assuming a binding affinity of $0.1\mu\text{M}$ |
| $k_{k1}, k_{k2}$ | $36 \text{ hr}^{-1}$ | Turnover rate for RsbW2 kinase: $10^{-3}$-$10^{-2} \text{ s}^{-1}$ [18] |
| $k_p$ | $180 \text{ hr}^{-1}$ | [15] |
| $k_{deg}$ | $0.7 \text{ hr}^{-1}$ | Based on $\sim 1\text{hr}$ doubling time |
| $v_0$ | $0.4 \mu\text{Mhr}^{-1}$ | Chosen to ensure total $\sigma^B$ level $\sim 1\mu\text{M}$ in the absence of stress [18] |
| $f$ | 30 | [13] |
| $K$ | $0.2 \mu\text{M}$ | Intermediate binding constant for σ-factor promoters (Typical range: $10^{-9}$-$10^{-6} \text{ M}$; [16]) |
| $\lambda_W, \lambda_V$ | 4, 4.5 | Varied |
| $[p_B]_{tot}$ | $0.05 \mu\text{M}$ | Assuming $\sim 50$ specific binding sites for $\sigma^B$ per genome |
| $RNApol_{tot}$ | $10 \mu\text{M}$ | [16] |

## Supplementary Information

### Supplementary Figures

**Figure S1.**
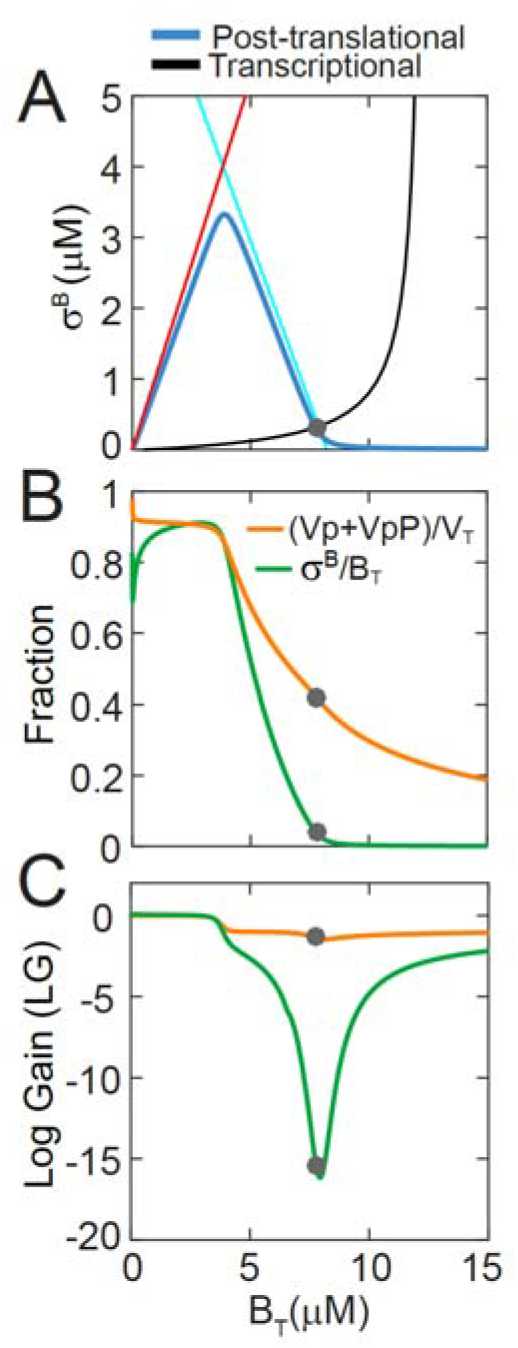
Ultrasensitive negative feedback in the σ^B^ network. **A.** Decoupled post-translational (blue curve) and transcriptional (black curve) responses of the σ^B^ network for *λ_W_* = *RsbW_T_*/*B_T_* = 4, *λ_V_* = *RsbV_T_*/*B_T_* = 4.5. σ^B^ and B_T_ represent the concentrations of free and total σ^B^. Gray circle marks the steady states of the full system. Red and blue lines represent the piecewise analytical approximations of the post-translational response. **B**. Decrease in the fraction of phosphorylated RsbV (V_p_+V_p_P – orange curve) and unbound (green curve) as a function total operon expression level according to the post-translational response. **C**. Sensitivity of the post-translational response of phosphorylated RsbV (Vp+VpP – orange curve) and unbound (green curve) to changes in total operon expression level (B_T_). At the shown steady state (gray circles) both responses have LG<-1.

**Figure S2.**
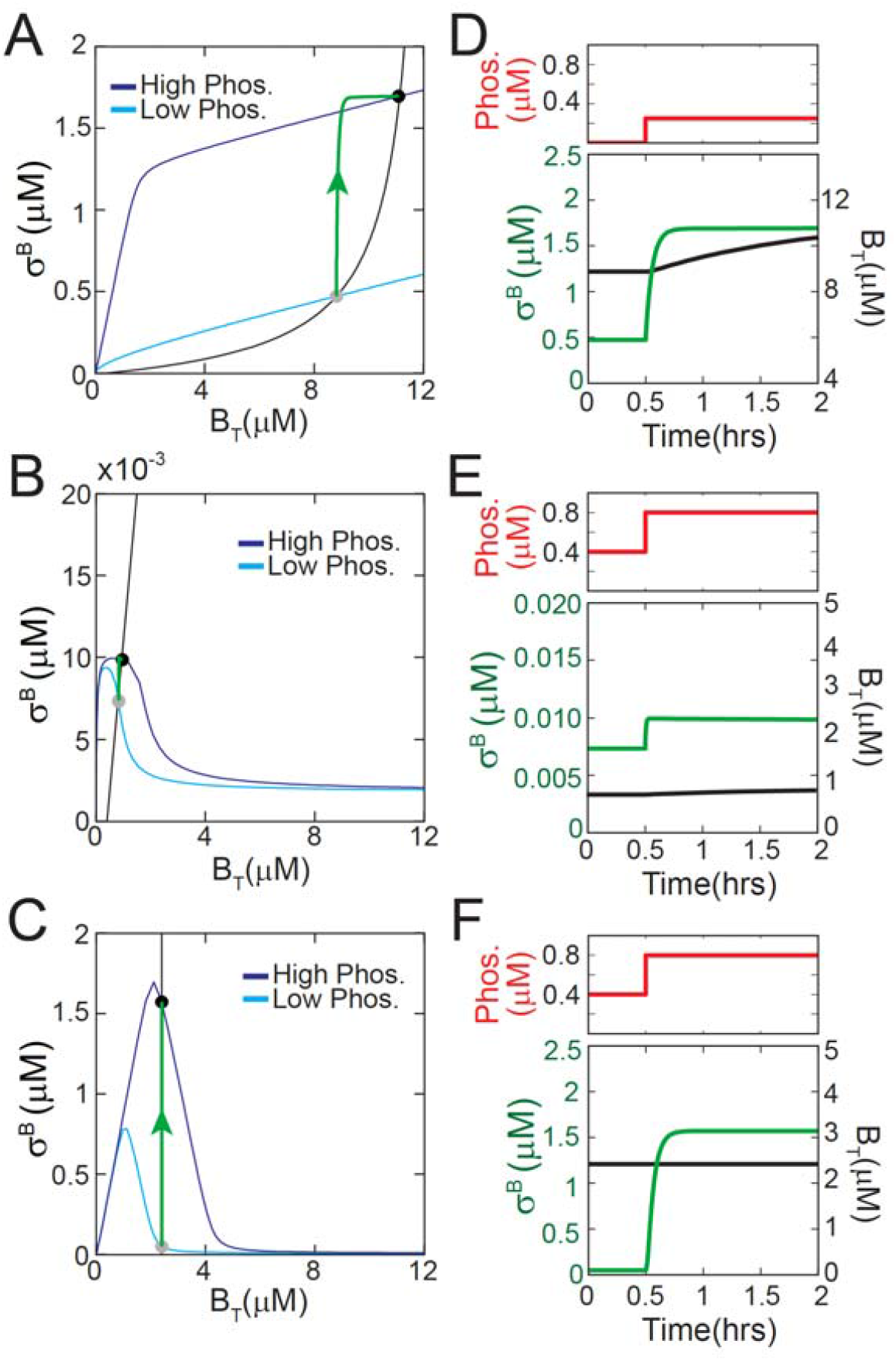
σ^B^ does not pulse for networks that lack negative feedback. **A-C.** Decoupled post-translational and transcriptional responses of σ^B^ networks that lack negative feedback. (A) λ_W_ = 2, λ_V_ = 2 (Region I in Fig. 2D) - positive feedback system; (B) λ_W_ = 8, λ_V_ = 4.5 (Region III in Fig. 2D) a non-responsive system; (C) λ_W_ = 4, λ_V_ = 4.5 with no transcriptional feedback – no feedback system. In each panel cyan and blue curves show the post-translational response at low and high phosphatase concentrations, and black curve shows the transcriptional response. Gray and black circles mark the steady states of the full system. Step-increase in phosphatase causes a shift in the post-translational response from low phosphatase-cyan to high phosphatase-blue and leads to an increase in σ^B^ (green curve) in all three systems. **D-F**. Time-course representations of green trajectories described in A-C. Note that σ^B^ does not pulse in any of the three systems.

**Figure S3.**
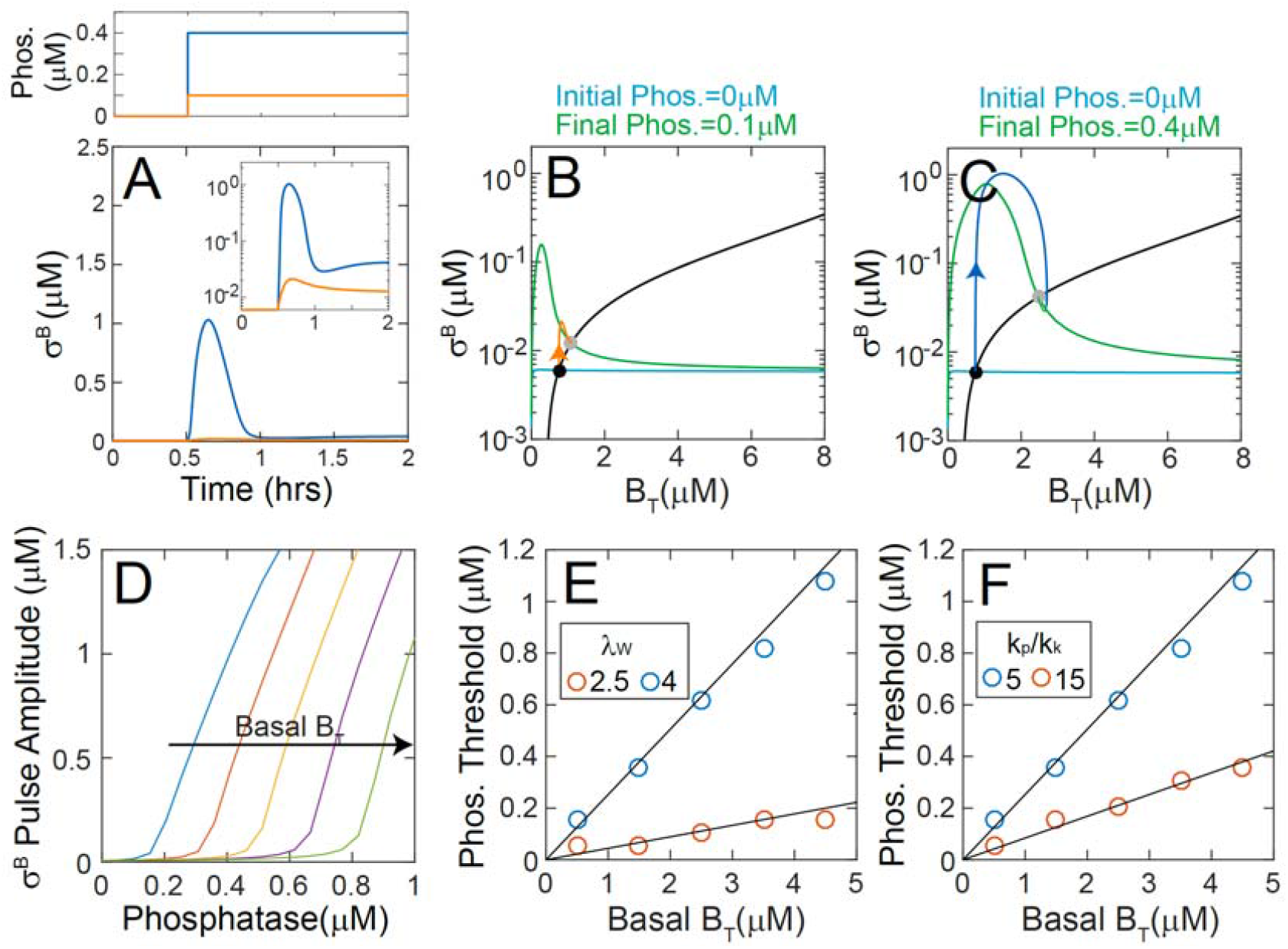
Dependence of σ^B^ pulse amplitude on phosphatase concentrations and post-translational parameters. **A.** Time-course representations of σ^B^ pulse trajectories for small (0.1μM-orange curve) and large (0.4μM-blue curve) step-increases in phosphatase. λ_W_ = 4, λ_V_ = 4.5 for both trajectories. **B,C**. Representation of the σ^B^ pulse trajectories and decoupled post-translational and transcriptional responses of σ^B^ network for small (B) and large (C) step-increases in phosphatase. Cyan and green curves show the post-translational responses at initial and final phosphatase levels. Black curves show the transcriptional response. Black and gray circles mark the steady states of the full system. Note that at the initial phosphatase level the σ^B^~0 and B_T_ is at the basal level of σ^B^ operon transcription. The small step-increase in phosphatase does not significantly shift the post-translational response around the initial steady state leading to minor, transient increase in σ^B^ (orange curve in B). The large step-increase in phosphatase (C) does significantly shift the post-translational response around the initial steady state leading to prominent pulse in σ^B^ (blue curve in C). D. σ^B^ pulse amplitudes show a threshold linear response to increase in phosphatase level. The threshold phosphatase level increases with increasing basal level of σ^B^ operon transcription (Basal B_T_). **E,F**. Phosphatase threshold for pulsing as a function of Basal B_T_ for different values of the (E) RsbW relative synthesis rate (λ_W_) and (F) the ratio of phosphatase to kinase rates (k_p_/k_k_). The circles represent threshold levels calculated from simulations. The black lines represent the analytical approximation: P_T_=V_0_*k_k_*(λ_W_/2 − 1 − λ_W_k_deg_/k_k_)/k_p_/k_deg_, where v_0_ and k_deg_ are the basal rate of σ^B^ operon transcription and protein degradation/dilution rate respectively. Basal B_T_ = v_0_ /k_deg_.

**Figure S4.**
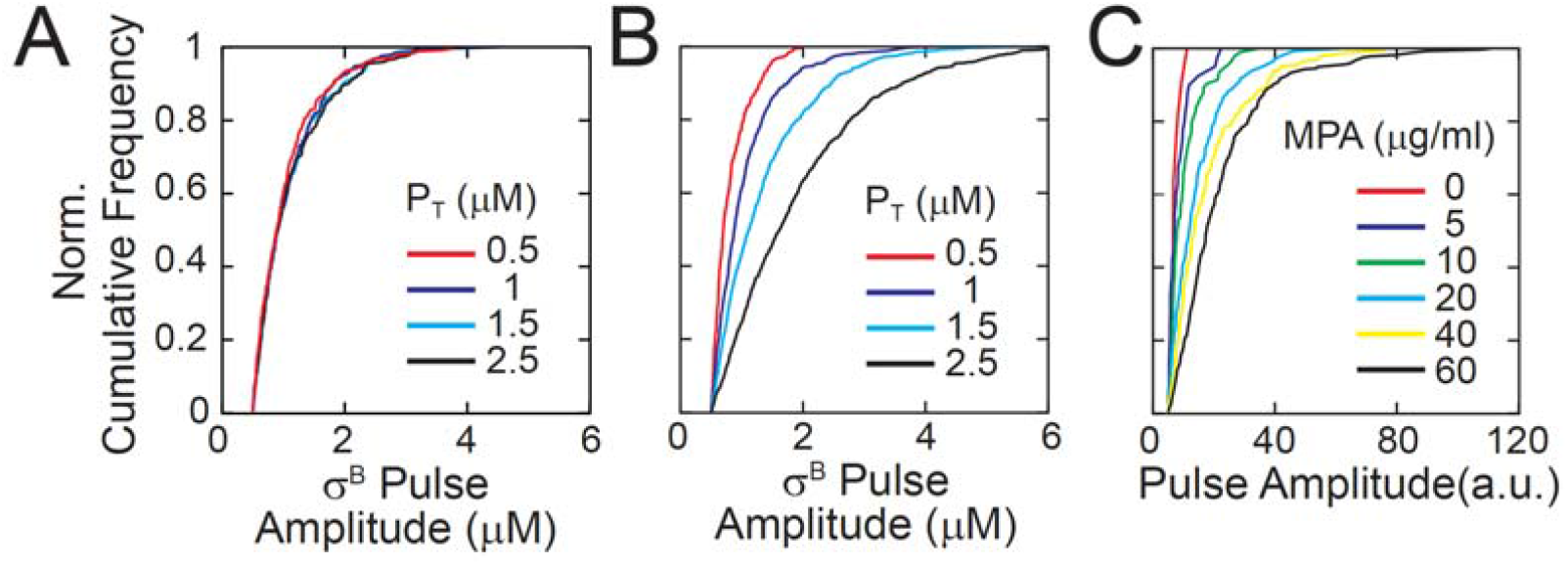
Pulsatile response of the σ^B^ network encodes phosphatase burst size not burst frequency. **A-C.** Simulation results for the response of the σ^B^ network model to stochastic fluctuations in levels of stress-sensing phosphatase RsbQP for fixed mean burst size and varying burst frequency. Green circles and errorbars show mean levels and standard deviations calculated from stochastic simulations. Black line is a linear fit. Mean σ^B^ pulse amplitude (A) is insensitive to mean phosphatase level. Mean σ^B^ pulse frequency (B) increases linearly as a function of mean phosphatase level. Mean σ^B^ target expression (C) increases non-linearly as a function of mean phosphatase level. **D-F.** Normalized pulse amplitude cumulative histograms for stochastic simulations with (D) burst frequency modulation, (E) burst-size modulation and (F) experimental data taken from [13]. Different colors represent varying levels of mean phosphatase (P_T_) in the model or mycophenolic acid (MPA, energy stress) in experiments.

**Figure S5.**
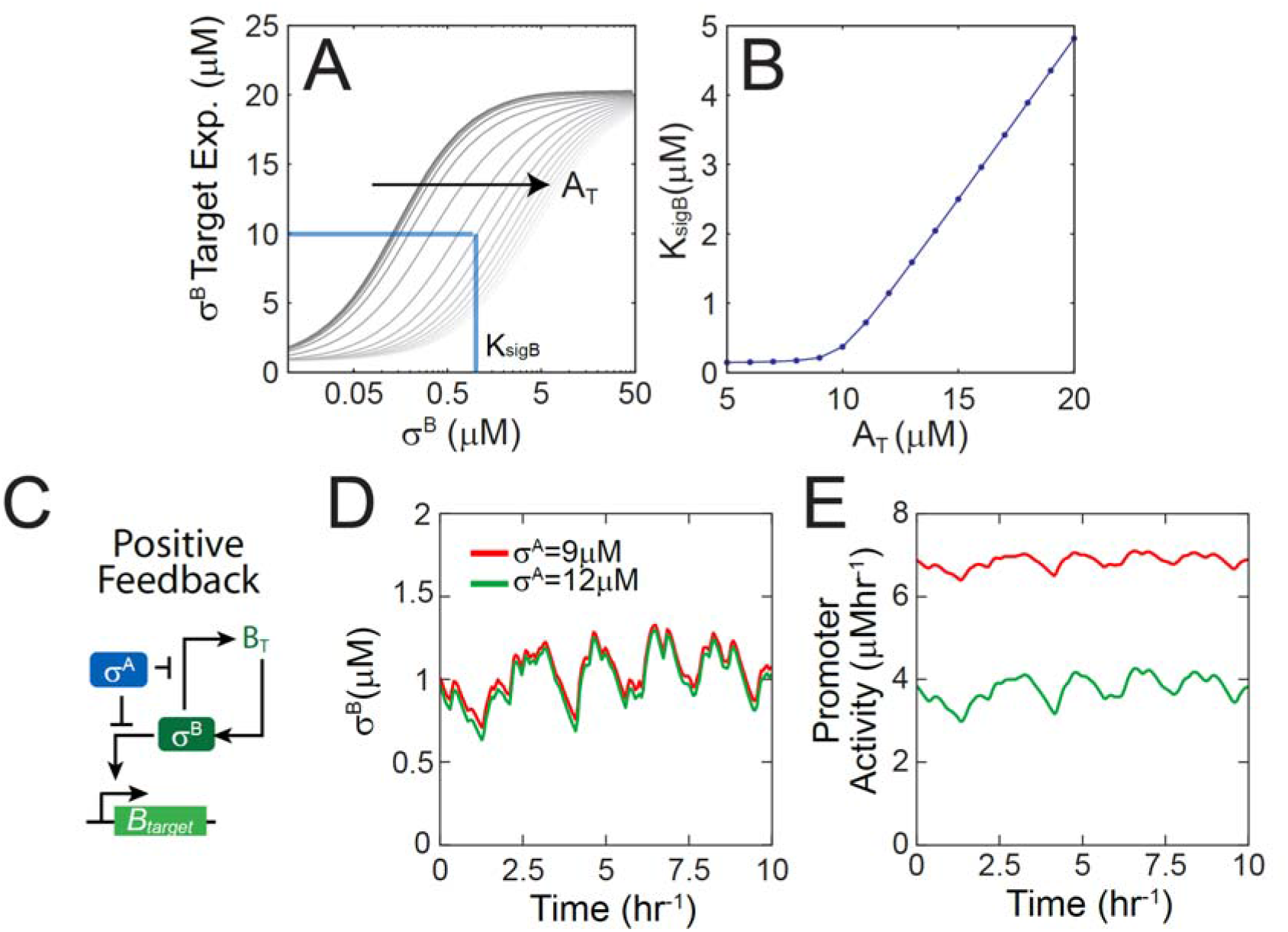
Sensitivity of the σ^B^ target expression to σ^A^ and competition for RNA polymerase. **A.** Steady-state dependence of the σ^B^ target expression on free σ^B^ for different total levels of the housekeeping σ-factor (A_T_). **B**. K_sigB_, the half-maximal constant of the dependence of σ^B^ target expression, as a function of the total levels of the housekeeping σ-factor (A_T_). **C**. Simplified network diagrams of a positive feedback regulated stress σ-factor σ^B^ competing with housekeeping σ-factor σ^A^ for RNA polymerase. **D-E.** Trajectories of free σ^B^ (D) and σ^B^ target promoter activity (E) in response to stochastic phosphatase input at two different levels of total σ^A^ (A_T_ = 9μM-low competition for RNA polymerase; A_T_ = 12μM-high competition for RNA polymerase).

**Figure S6.**
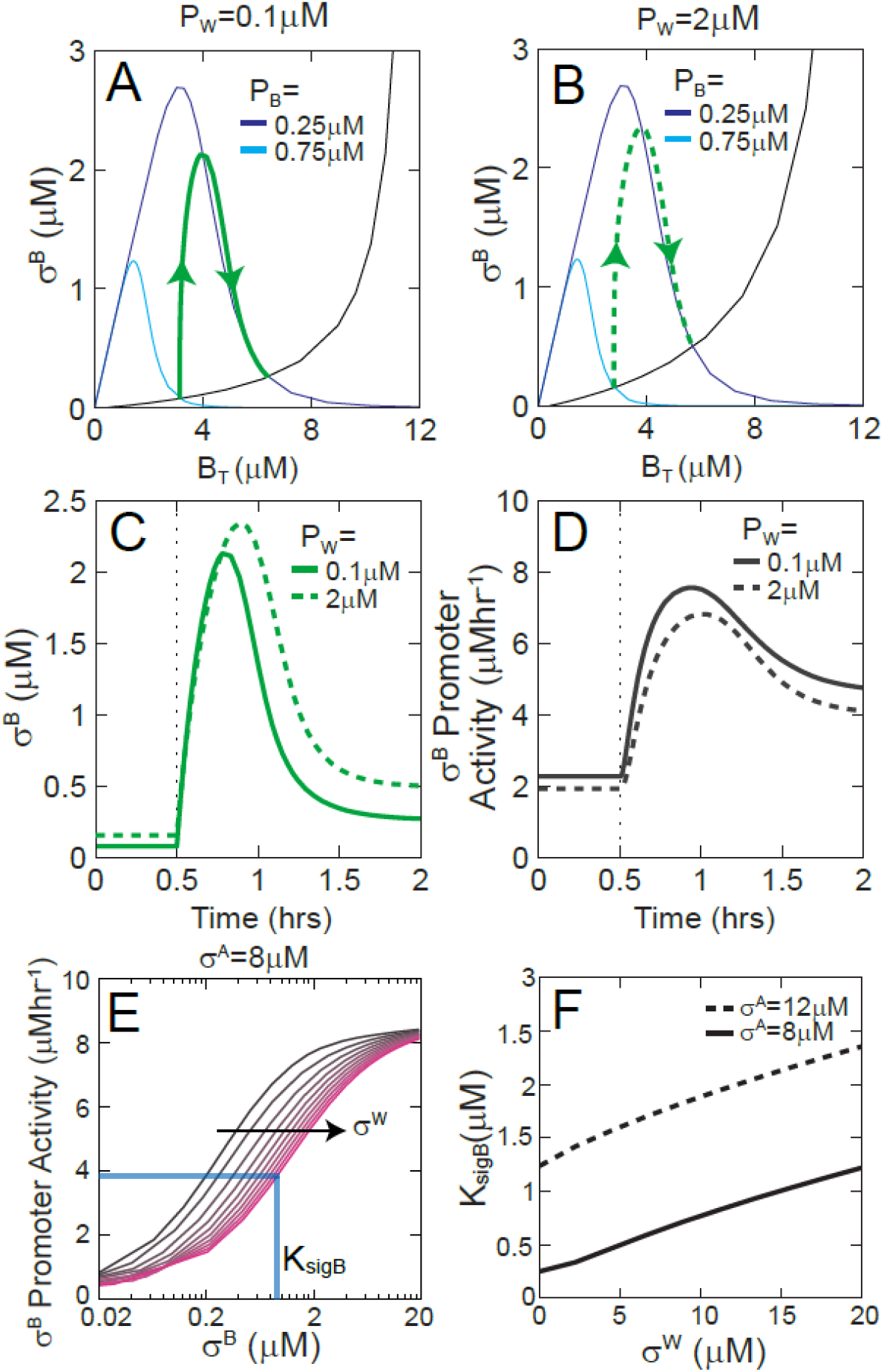
Pulsatile response and RNApol competition in the multiple stress σ-factor model. **A,B**. Decoupled σ^B^ post-translational and transcriptional components in the simplified model for the competition of stress σ-factors. Cyan and blue curves show the post-translational response at low and high concentration of σ^B^ stress signaling protein P_B_. Black curve shows the transcriptional responses. A step-increase in P_B_ causes a shift in the steady-state post-translational response (from low phosphatase-cyan to high phosphatase-blue) and leads to a pulsatile σ^B^ response trajectory (green curve). Concentration of σ^W^ stress signaling protein P_W_ was kept fixed at 0.1 μM (A) and 2 μM (B). **C,D**. Time-course representations of the green trajectories in (A,B) showing σ^B^ (C) and σ^B^ promoter activity (D) respectively. **E**. Steady state dependence of the concentration of σ^B^ target promoter activity, on the level of free σ^B^ for different levels of the σ-factor σ^W^. **F**. K_sigB_, the half-maximal constant of the dependence of target expression on σ^B^ as a function of the concentration of the stress σ-factor σ^W^ for different levels of the housekeeping σ-factor σ^A^. Total RNA polymerase core concentration was kept fixed at 10μM for all simulations.

## Supplementary Methods

### *Derivation of steady state asymptotes for the* σ^B^ *post-translational response*

To understand the steady state post-translational response of σ^B^ network (Figs. 1, 2) we used the mass balance for the operon components RsbW, RsbV and σ^B^ together with the phosphate flux balance to derive approximate dependence of free σ^B^ on B_T_. We found that the post-translational response of the network varies depending on whether the concentration of operon components is lower or higher than a threshold level defined by the concentration of the stress phosphatase P_T_.

For low B_T_ (B_T_ < 2P_T_k_p_/k_k_/min[λ_W_, λ_V_]), the maximum phosphatase flux (k_P_*P_T_) exceeds the maximum kinase flux (k_k_*min[W_T_, V_T_]/2) and as a result, V_P_≈0. In addition most of the anti-anti-σ-factor V is in the W_2_V_2_ complex. Taking this into account and applying the mass balance for RsbV,

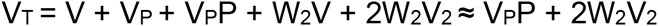

Next applying the balance for kinase and phosphatase fluxes,

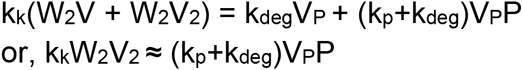

This implies that: W_2_V_2_ = min[V_T_/(2 + k_k_/(k_p_+k_deg_)),W_T_/2], where the minimum function is applied to account for the fact that the concentration of W_2_V_2_ cannot exceed half the total RsbW concentration.

Using the above equation in the mass balance for W we solve for W2B and thereby σ^B^,

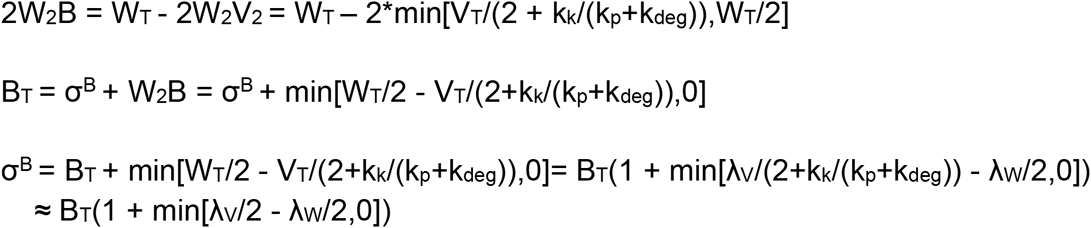

In contrast, for higher B_T_ (B_T_>2P_T_k_p_/k_k_/min(λ_W_, λ_V_)), where the RsbW kinase dominates the phosphatase, V_P_ is not negligible and the phosphatase is saturated (V_P_P≈P_T_). Again using this in the mass balance for RsbV,

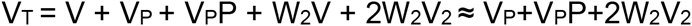

And applying the balance for kinase and phosphatase fluxes,

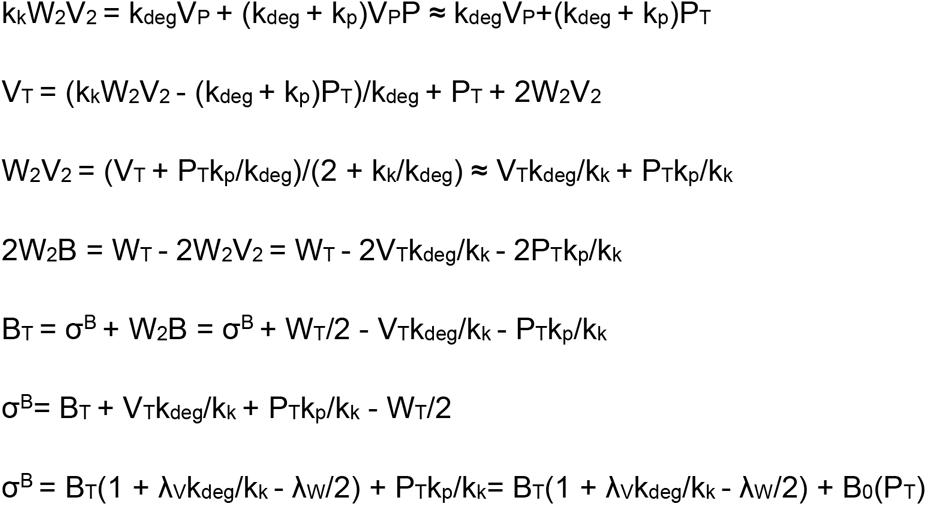

where B_0_(P_T_) = P_T_k_P_/k_k_

Note that since σ^B^ concentration cannot be negative this approximation only applies for B_T_<B_0_(P_T_)/(λ_W_/2 − 1 − λ_W_k_deg_/k_k_). For higher B_T_, σ^B^ ~0.

Taken together the dependence of σ^B^ on B_T_ can be described by the following system of equations:

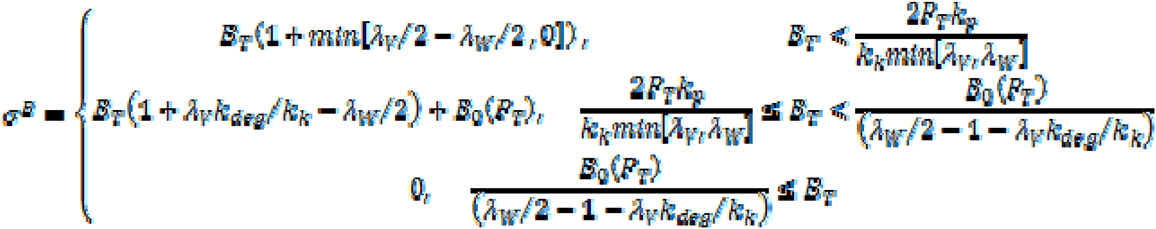

Based on the above equation, the sensitivity of the σ^B^ post-translational response depends on (λ_W_, λ_V_), i.e. the stoichiometry of operon components:

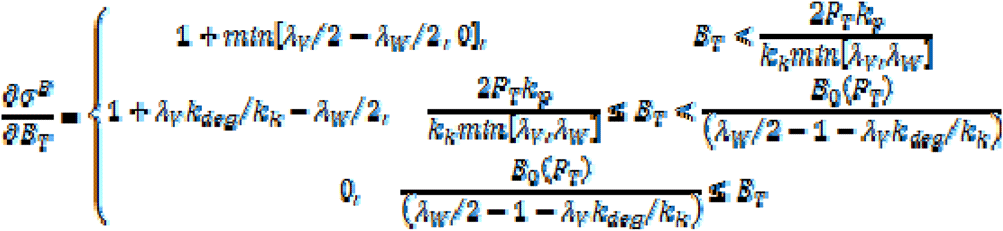

Which shows that the (λ_W_, λ_V_) parameter space can be divided into three regions based on qualitative differences in the post-translational response.

**Region I (λ_W_<2+λ_V_k_deg_/k_k_):** *∂*σ^B^/*∂*B_T_>0 and free σ^B^ increases as a function of B_T_ irrespective of P_T_.

**Region II (2+2*λ_V_k_deg_/k_k_<λ_W_<2+λ_V_):** *∂*σ^B^/*∂*B_T_>0 for B_T_<2P_T_k_p_/k_k_/min[λ_W_, λ_V_] and *∂*σ^B^/*∂*B_T_<0 for B_T_>2P_T_k_p_/k_k_/min[λ_W_, λ_V_]. Thus free σ^B^ is a non-monotonic function of B_T_.

**Region III (λ_W_>2+λ_V_):** *∂*σ^B^/*∂*B_T_<0 and free σ^B^ decreases as a function of B_T_ irrespective of P_T_.

Thus the asymptotic description shows how relative synthesis rate of σ^B^ operon partners by controls the sign of post-translational response sensitivity (*∂*σ^B^/*∂*B_T_). Specifically it shows that *∂*σ^B^/*∂*B_T_ <0 is only possible in Region II where 2 + 2*λ_V_k_deg_/k_k_ < λ_W_ < λ_V_ + 2. This implies that the overall feedback in the σ^B^ network can only be negative in Region II, thereby explaining why pulsatile responses are only seen combinations sampled from this region (Fig. 1B-D). Note also that the boundary equations for Region II closely approximate the boundaries of the operon stoichiometry space calculated by sampling (λ_W_, λ_V_) combinations (Fig. 2D).

The asymptotic description also explains (Fig. S3) the observation that a threshold level of phosphatase is essential for pulsing [13]. As shown above, *∂*σ^B^/∂B_T_<0 in Region II only when:

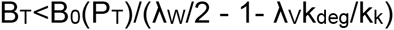

where B_0_(P_T_) = P_T_k_p_/k_k_.

However B_T_>v_0_/k_deg_ since σ^B^ operon transcription has a basal rate independent of σ^B^ level. v_0_ and k_deg_ are the basal rate of transcription and protein degradation/dilution rate respectively.

Consequently, *∂*σ^B^/*∂*B_T_<0 in Region II only for:

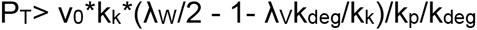

This defines the threshold level of phosphatase essential for *∂*σ^B^/*∂*Bτ<0 and for the σ^B^ network to operate in a negative feedback regime. As a result, the σ^B^ network only pulses for phosphatase levels above this threshold. Note that this threshold level is proportional to both the basal level of σ^B^ operon expression and the ratio of kinase to phosphatase rate constants and increases as a function of the RsbW synthesis ratio λ_W_ (Fig. S3). This indicates that it represents the basal level of kinase flux that the stress-regulated phosphatase flux must exceed to trigger a response.

### Mathematical model of competition between σ^B^ and σ^A^

#### *Additional reactions for the model of competition between* σ^B^ *and* σ^A^

To model the competition for RNA polymerase between σ^B^ and the housekeeping σ-factor σ^A^ (Figs. 5 and S5), we extended the model described above and supplemented reactions (1–9) with the following reactions:

- Reversible binding of σ-factors and RNA polymerase

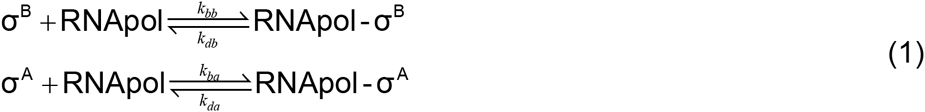
- Reversible binding of RNApol-σ^B^ complexes to target promoters

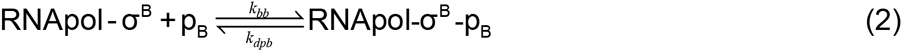
- Production of σ^B^, RsbW and RsbV

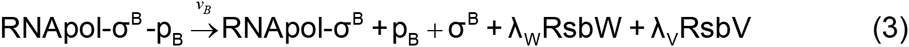

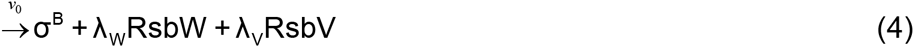

Where v_0_ is the basal synthesis rate and v_B_=v_0_*f/[p_B_]_tot_ is the maximal rate. *f* is the fold change in protein synthesis due to positive autoregulation and [p_B_]_tot_ is the total concentration of the σ^B^ promoter.

#### Model equations

The following set of equations that describe network dynamics of this extended model:

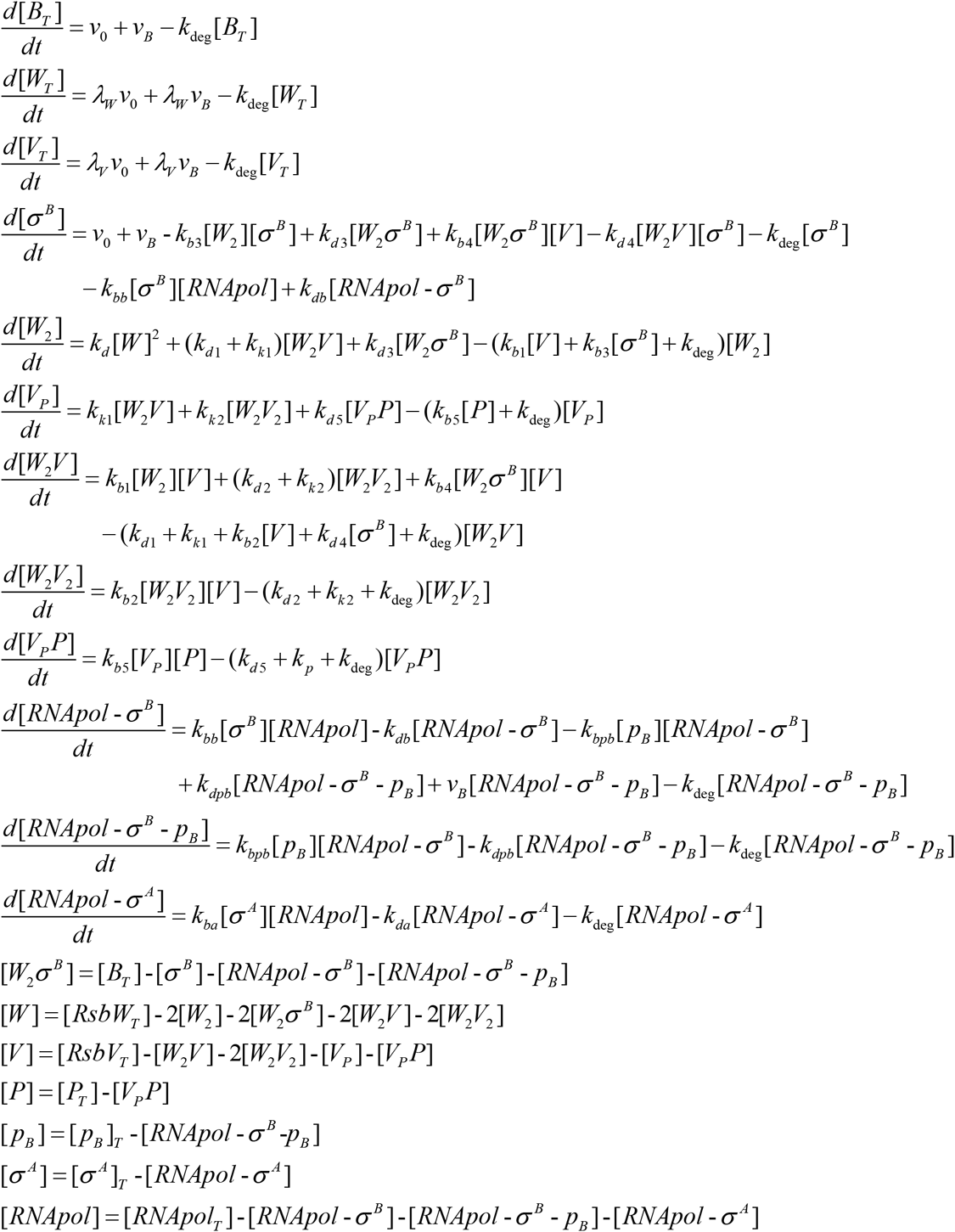

#### *Model equations for the model of competition between* σ^B^, σ^W^ *and* σ^A^

To model the competition for RNA polymerase between σ^B^ the housekeeping σ-factor σ^A^ and the alkaline stress response σ-factor σ^W^ (Figs. 6 and S6), we simplified the model for the post-translational control of stress σ-factors while explicitly including reactions for the binding/unbinding of RNA polymerase, σ-factors and target promoters. This model included the following set of equations:

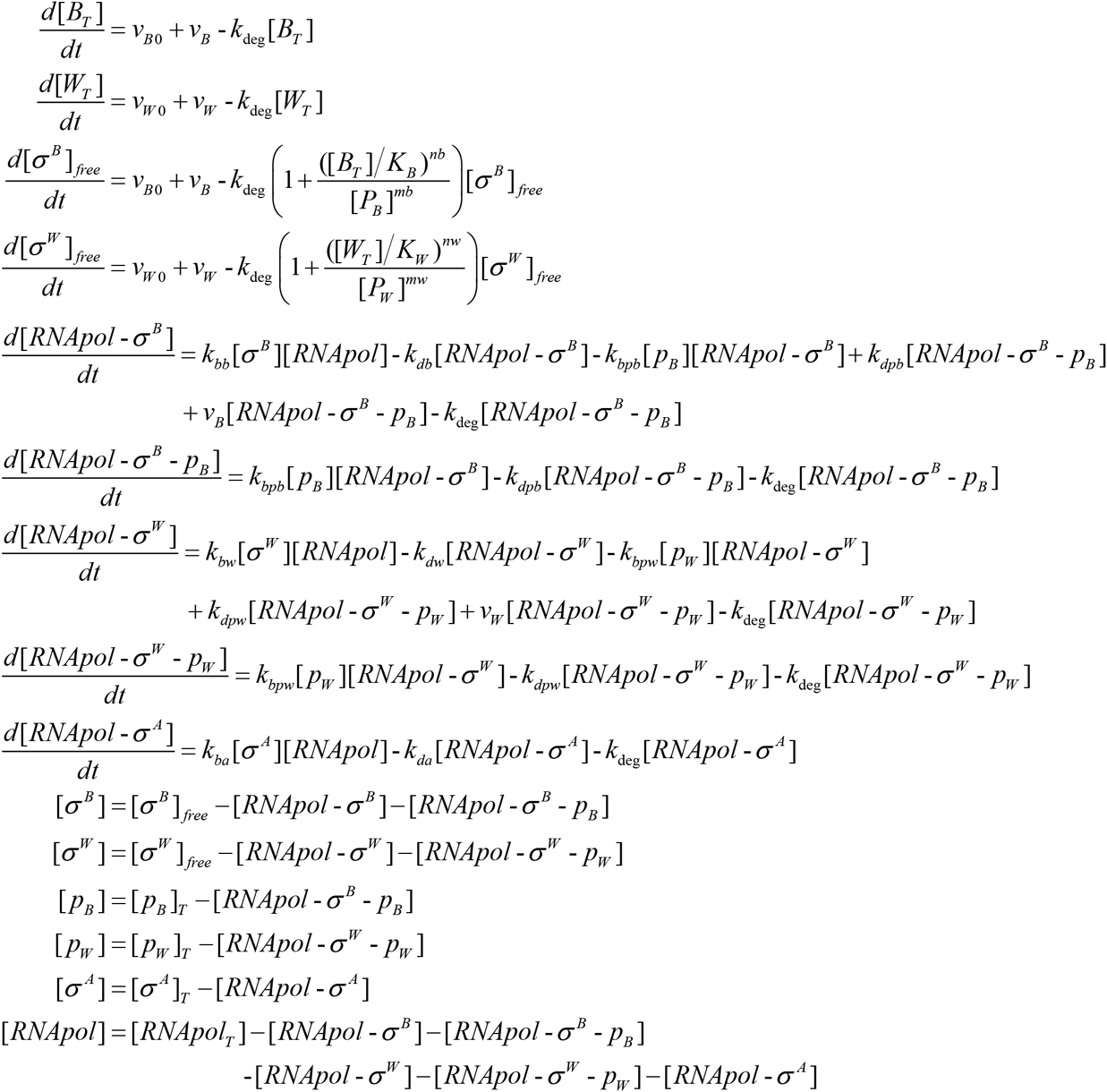

